# Subjective confidence reflects representation of Bayesian probability in cortex

**DOI:** 10.1101/2021.04.10.439272

**Authors:** Laura S. Geurts, James R. H. Cooke, Ruben S. van Bergen, Janneke F. M. Jehee

## Abstract

What gives rise to the human sense of confidence? Here, we tested the Bayesian hypothesis that confidence is based on a probability distribution represented in neural population activity. We implemented several computational models of confidence, and tested their predictions using psychophysics and fMRI. Using a generative model-based fMRI decoding approach, we extracted probability distributions from neural population activity in human visual cortex. We found that subjective confidence tracks the shape of the decoded distribution. That is, when sensory evidence was more precise, as indicated by the decoded distribution, observers reported higher levels of confidence. We furthermore found that neural activity in the insula, anterior cingulate, and prefrontal cortex was linked to both the shape of the decoded distribution and reported confidence, in ways consistent with the Bayesian model. Altogether, our findings support recent statistical theories of confidence and suggest that probabilistic information guides the computation of one’s sense of confidence.

## Introduction

Virtually any decision comes with a sense of confidence – a subjective feeling that clearly affects our everyday choices. For example, we reduce speed when driving at night because we feel less confident about our estimates of distance to surrounding traffic, we hesitate to try a piece of food when unsure about its taste, and resist investing in stocks unless convinced of their likely future profit. But what is this sense of confidence that accompanies most all of our decisions?

Recent Bayesian decision theories^1–5^ propose that confidence corresponds to the degree of belief, or probability, that a choice is correct based on the evidence. More specifically, these theories propose that confidence is a function of the posterior probability of being correct, which links confidence directly to the quality of the evidence on which the decision is based. Thus, greater imprecision in evidence reduces the probability that the choice is correct, which should result in lower levels of confidence. The agent’s evidence is similarly described as a degree of belief in an event, or more formally, as a probability distribution over a latent variable. For example, the evidence could be a probability distribution over perceived distance to surrounding traffic. The width of the distribution (range of probable distances) is broader in the dark than on a clear day, thereby signaling greater imprecision or uncertainty. Although central to the Bayesian confidence hypothesis, whether such probabilistic representations play a role in confidence is currently unclear.

Results from behavioral studies^6–9^ are consistent with the notion that confidence is computed from the degree of imprecision in sensory information. However, a major limitation of this work has been the use of physical sources of noise, such as a variation in image brightness or contrast, to manipulate uncertainty. This is problematic because it could be that observers simply monitor such stimulus properties as external cues to uncertainty and confidence^7, 10–13^. While physiological studies have found neural correlates of statistical confidence in the orbitofrontal^14, 15^ and lateral intraparietal cortex^16^, these studies used a two-alternative forced choice (2AFC) task, so that the animal could simply rely on the distance between stimulus estimates (i.e. point estimates) and category boundary to compute confidence^17^, and need not use a representation of probability. Thus, one of the most fundamental assumptions of normative theories of decision-making – that confidence is derived from a probabilistic representation of information – has yet to be tested in cortex.

Here, we use a combination of functional Magnetic Resonance Imaging (fMRI), psychophysics, and computational modeling to address two fundamental questions. 1) Is confidence based on a probabilistic representation of sensory information? And if so, 2) what neural mechanisms extract confidence from this cortical representation of uncertainty? Human participants viewed random orientation stimuli, and reported both the orientation of the stimulus and their level of confidence in this judgment. Critically, no physical noise was added to the stimuli. We quantified the degree of uncertainty associated with stimulus representations in visual cortex using a probabilistic decoding approach^10, 18^, relying on trial-by-trial fluctuations in internal noise to render the evidence more or less reliable to the observer. We used the decoded probability distributions to compare between human data and simulated data from a Bayesian observer, as well as two alternative models implementing heuristic strategies to confidence. Corroborating the Bayesian model, we discovered that human confidence judgments track the degree of uncertainty contained in visual cortical activity. That is, when the cortical representation of the stimulus was more precise (as indicated by a narrower decoded probability distribution), participants reported higher levels of confidence. In addition, activity in the dorsal Anterior Insula (dAI), dorsal Anterior Cingulate (dACC) and rostrolateral Prefrontal Cortex (rlPFC) reflected both this sensory uncertainty and reported confidence, in ways predicted by the Bayesian observer model. Taken together, these results support normative theories of decision-making, and suggest that probabilistic sensory information guides the computation of one’s sense of confidence.

## Results

### Ideal observer models

The observer’s task is to infer the orientation of a stimulus from a noisy sensory measurement, and report both this estimate and their level of confidence in this judgment. We consider three model observers for this task. The decision process is identical for all three observers, but they use different strategies to confidence.

The observer’s measurement *m* of the sensory stimulus *s* is corrupted by noise: even when the physical stimulus is held constant, the measurement varies from trial to trial. Thus, the relationship between stimulus and measurement on each trial is given by a probability distribution, *p*(*m*|*s*) which we model as a circular Gaussian centered on the stimulus and with variance *σ*_*m*_^2^(*s*). This variability in the measurements stems from various sources of noise that are of both sensory and non-sensory origin. Specifically, we consider three sources of noise: two sensory and one non-sensory. The first source depends on stimulus orientation, with larger noise levels for oblique than cardinal stimulus orientations. This pattern captures the well-established ‘oblique effect’ in orientation perception^19, 20^. The second source varies in magnitude from trial to trial, and captures, for example, random fluctuations in neural response gain in sensory areas^21^. Finally, non-sensory noise refers to those sources of variance that affect, for example, the stimulus representation while held in working memory, or task-related processes in areas downstream of sensory cortex.

To infer the orientation of the stimulus from the measurement, all three observers invert the generative model to compute the posterior probability distribution *p*(*s*|*m*) (Equation 12). This distribution quantifies the degree to which different stimulus values are consistent with the measurement. The mean of the posterior distribution is the model observer’s estimate of the stimulus *ŝ*. We take the (circular) variance of the distribution as a measure of the degree of uncertainty in this estimate. The observer’s internal estimate of orientation is subsequently translated into an overt (behavioral) response, *r*. This transformation from internal estimate into motor response is noisy. Thus, across trials, the response fluctuates around *ŝ*, where (motor) noise is drawn from a circular Gaussian (Equation 13).

How does each of the observer models compute confidence? The ideal strategy is to consider the degree of imprecision in the observer’s decision, which depends on all sources of variance that affect their reports. Specifically, for the estimation task used here, it is statistically reasonable to compute confidence as a function of the expected magnitude of the error in the observer’s response. We quantified this as follows:

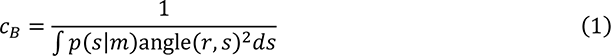

where *c*_*B*_ refers to the reported level of confidence, and angle(*r*, *s*)^2^ represents the magnitude of the response error (i.e., the squared acute-angle distance between response and (latent) stimulus). In words, when uncertainty in evidence is higher, the expected decision error tends to be larger, and reported confidence will be lower. However, our predictions do not strongly depend on the particular function assumed here, as long as confidence monotonically decreases when overall uncertainty increases. We refer to this model as the Bayesian or Probabilistic observer, as confidence is based (in part) on the posterior probability distribution – a probabilistic notion of uncertainty.

The second model observer uses certain properties of the stimulus, such as its orientation, as a cue to confidence. This observer has learned through experience that behavioral precision is usually better for cardinal than for oblique orientations. The observer utilizes this learned relationship as a heuristic, and simply reports lower levels of confidence for those orientations that generally result in reduced levels of performance. We refer to this model as the Stimulus heuristics observer, and formally define their confidence as:

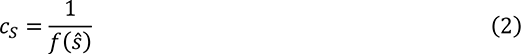

where *f*(*ŝ*) is a function that rises for oblique orientations (see Equation 14 in Methods). As the strategy ignores many sources of noise that create uncertainty, it is clearly suboptimal, but it could potentially explain human behavior, which is why we include the strategy here.

The third and final model observer ignores the imprecision in internal estimates altogether, and computes confidence exclusively from the noise in their motor response. We refer to this model as the Response heuristics observer. That is, on a given trial the observer simply notices a large offset between their internal orientation estimate and overt (motor) response. Observing that their response is off, they report lower levels of confidence. This is not an ideal strategy, but it is nonetheless a strategy that could result in a reliable link between confidence and behavioral performance, as we will show in our simulations below. We define confidence for this observer model as:

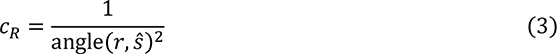

Where angle(*r*, *s*)^2^ is the squared acute-angle distance between orientation estimate *ŝ* and response *r*. Fig. 1 summarizes the three observer models.

**Fig. 1.**
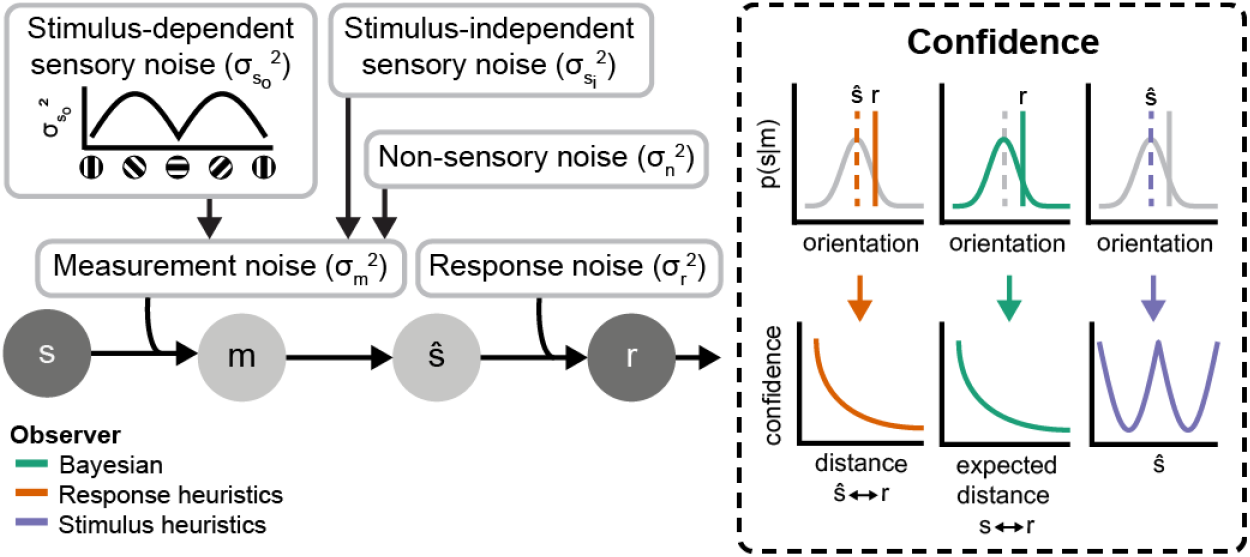
Overview of sources of noise and three observer models. The observers’ task is to estimate the presented stimulus orientation *s* from a noisy measurement m. Multiple sources of noise affect the perceptual decision-making process. The measurements (*m*) vary from trial to trial due to sensory sources of noise, which can be decomposed into stimulus-related (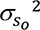) and stimulus-independent (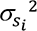) noise, as well as (unexplained) downstream (non-sensory) noise (*σ*_*n*_ ^2^). The observers compute their stimulus estimates ŝ as the mean of the posterior distribution *p*(*s*|*m*). The internal orientation estimate is transformed into a behavioral (overt) response *r*, which is subject to further noise (*σ*_*r*_ ^2^). The observer also gives their level of confidence in this behavioral estimate. The Bayesian observer computes confidence as a function of the expected distance between latent stimulus and response, which depends on both the response itself, and the width of the posterior *p*(*s*|*m*), which incorporates all sources of measurement noise. The Stimulus heuristics observer computes confidence as a function of their perceptual orientation estimate (*ŝ*). The Response heuristics observer computes confidence as a function of the distance between internal orientation estimate (*ŝ*) and overt motor response (*r*). Both Heuristics observers ignore the degree of uncertainty in their orientation estimates when computing confidence.

### Model predictions

What behavioral patterns should one observe for the different strategies to confidence? To address this question, we simulated the behavioral orientation estimates and associated confidence reports of the three model observers. As we will show below, this leads to a set of concrete predictions that we can then test in psychophysical and neuroimaging experiments. Does confidence predict behavioral performance? To address this question, we binned the simulated data according to reported level of confidence, and calculated the across-trial variance in behavioral orientation estimates for each of the bins. We first did this irrespective of the orientation of the stimulus. We found that the orientation judgments of the model observers were generally more precise when confidence was higher, regardless of the strategy to confidence employed by the observer (Fig. 2a). Thus, a predictive link between confidence and behavioral precision is consistent with several strategies, and does not necessarily imply that confidence is based on a probabilistic representation of the degree of uncertainty in one’s evidence.

**Fig. 2.**
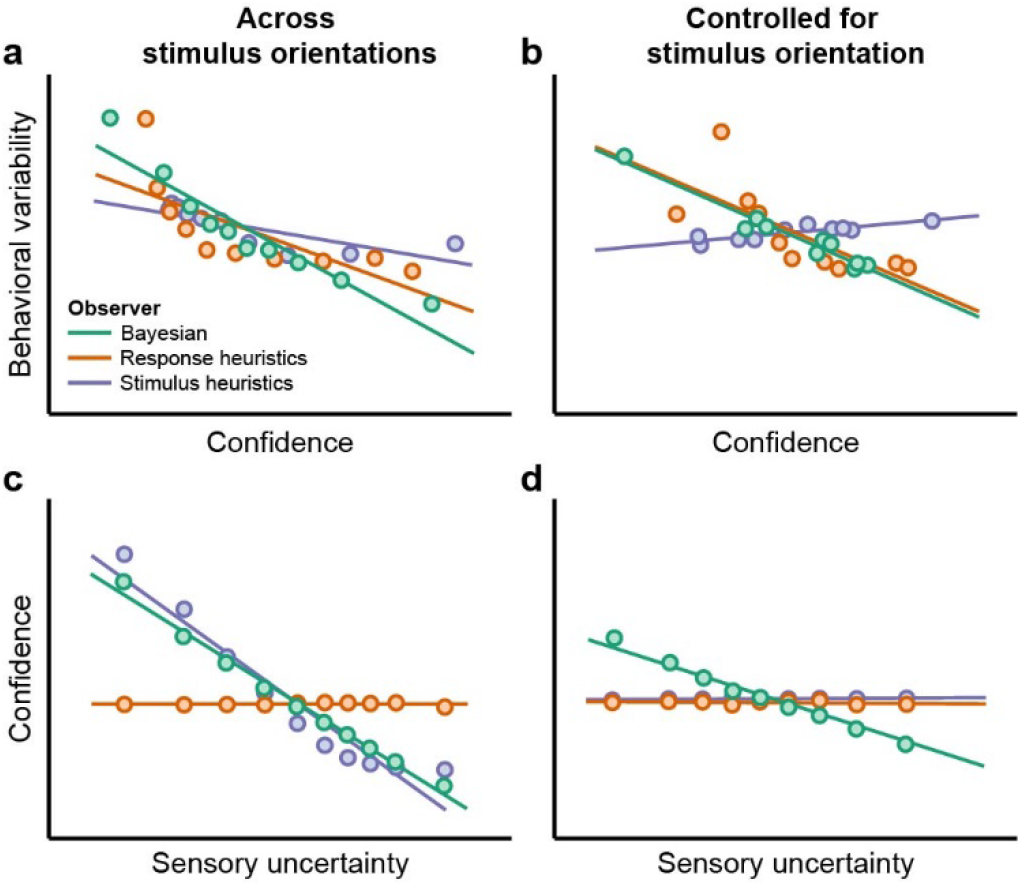
Ideal observer predictions. (a) Relationship between confidence and behavioral variability for a uniform stimulus distribution (orientation range: 0-179°). Trials (N = 50,000) were binned into ten equal-size bins of increasing confidence. For each bin, the variance of orientation estimation errors was computed and plotted against the mean level of confidence in that bin. Lines represent best linear fits. (b) Same as (a), but holding the stimulus constant. Confidence values were z-scored per observer such that they fall in the same range for all models. (c) Relationship between sensory uncertainty and confidence for a uniform stimulus distribution (orientation range: 0-179°). For visualization purposes, trials were binned into ten equal-sized bins of increasing uncertainty. The mean of both confidence and sensory uncertainty was computed across all trials in each bin, and is shown in the plot. Lines represent linear best fits computed on single-trial (unbinned) data. (d) Same as (c), but controlled for stimulus orientation.

We next turned to the relationship between confidence and behavioral performance for a constant stimulus. Closely replicating the experimental analysis procedures (see below), we first removed the effect of stimulus orientation from confidence, binned the data according to residual level of confidence, and calculated the variance in behavioral orientation estimates for each of the bins. We found that higher levels of confidence again predicted greater behavioral precision for both the Probabilistic and Response heuristics model (Fig. 2b). For the Stimulus heuristics observer, in contrast, we observed no clear link between confidence and behavioral performance. This makes sense, as this observer uses orientation as a cue to confidence, so an identical orientation stimulus should, when averaged across repeated presentations, always result in the same level of confidence, irrespective of any stimulus-independent sources of variance. Thus, this analysis could potentially enable us to differentiate between some, though not all, strategies to confidence.

We next considered the relationship between confidence and the quality of the observer’s evidence. Specifically, we determined the extent to which the degree of uncertainty in their sensory evidence predicted reported levels of confidence. Sensory uncertainty was quantified as the width of a probability distribution (see Methods), similar to the empirical conditions. For practical reasons, we here disregard the contribution of non-sensory sources of variance and focus on sensory uncertainty alone, so as to closely match the empirical analyses. Data were binned for visualization only, and mean levels of confidence and uncertainty were computed for each of the bins. When analyzed across stimulus orientation, and for both the Stimulus heuristics and Bayesian observer, reported levels of confidence consistently decreased as sensory uncertainty increased. However, we observed no such relationship between confidence and uncertainty for the Response heuristics observer (Fig. 2c). When holding the stimulus constant, the results were even more distinct between confidence strategies. That is, after we removed the contribution of stimulus orientation (see Methods), the relationship between sensory uncertainty and confidence still held for the Bayesian observer, but no such link between the fidelity of the observer’s sensory representation and confidence was observed for the two remaining models (Fig. 2d). This illustrates the importance of considering internal levels of uncertainty when studying confidence, and moreover indicates that these analyses, when combined, should enable us to adjudicate between strategies to confidence.

In sum, if human confidence estimates are based on probabilistic computations, then 1) behavioral variance in an orientation judgment task should be higher with reduced levels of confidence for a constant stimulus; 2) there should be an inverse relationship between sensory uncertainty in cortex and reported confidence; and 3) this inverse relationship should hold both across orientations and when holding the stimulus constant. With these predictions in hand, we now turn to the experimental data to see which strategy best describes human confidence judgments.

### Human observers

Do human observers use a probabilistic representation of evidence quality when reporting confidence? To address this question, we presented 32 human participants with oriented gratings while we measured their brain activity using fMRI. Observers reported the orientation of the grating, as well as their confidence in this judgment (see Extended Data Fig. 1). They generally performed well on this task, with a mean absolute behavioral estimation error of 4.34° ± 0.212° (mean ± SEM across subjects).

We first focused on the link between behavioral performance and confidence. For each observer, we divided all trials, regardless of presented orientation, into ten bins of increasing confidence, and computed and compared behavioral variability and mean level of confidence in each bin. We found a significant inverse relationship between confidence and behavioral variability (t(287) = −16.79, p < 0.001, r = −0.70, 95% CI = [-0.76, -0.64]; Fig. 3a, left). Thus, the observers’ orientation judgments were more precise when confidence was high. This indicates that the participants were able to meaningfully estimate their own level of confidence in the task.

**Fig. 3.**
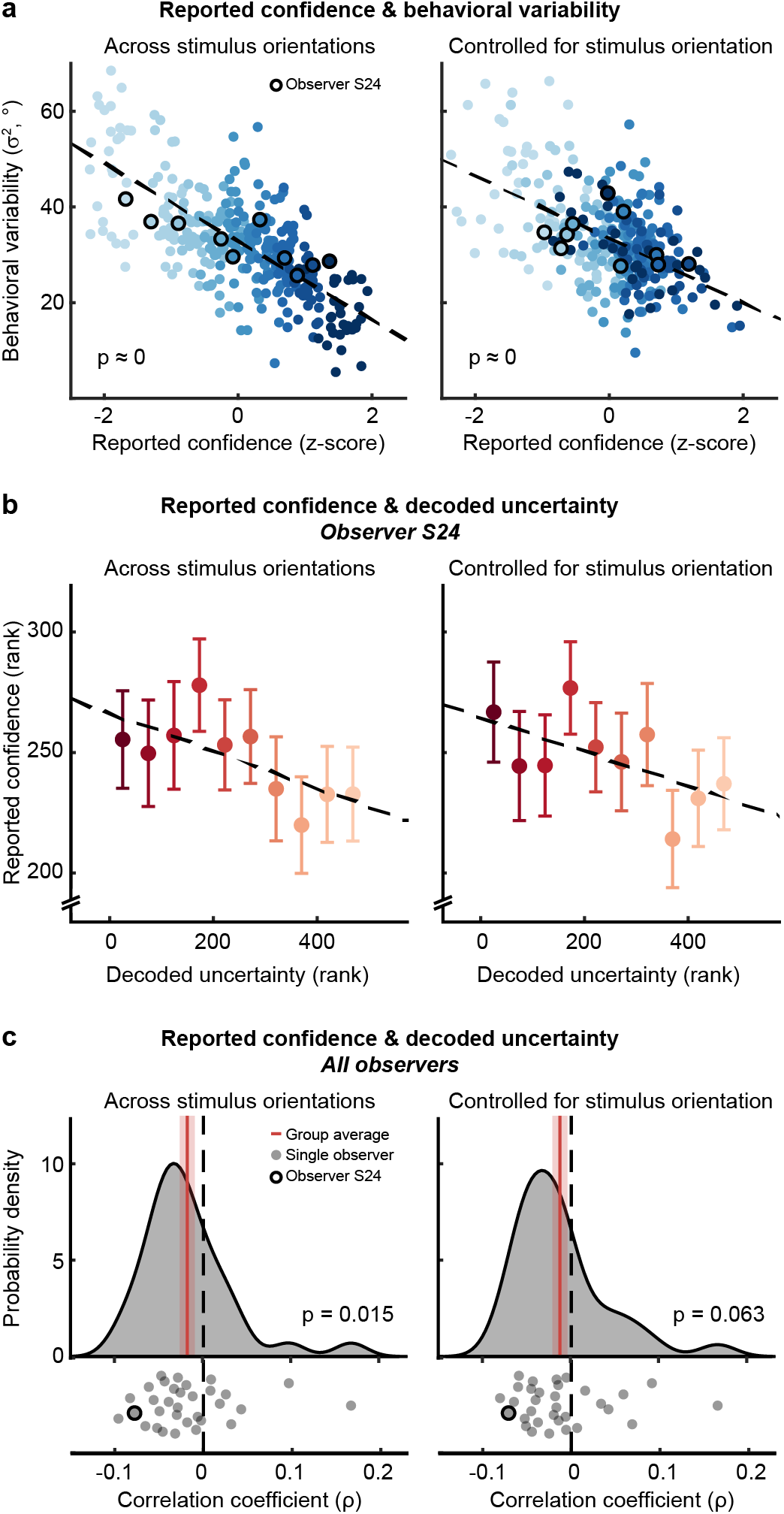
Reported confidence, behavioral variability, and decoded sensory uncertainty. (a) Behavioral variability decreases as confidence increases (left panel: t(287) = -16.79, p < 0.001, r = -0.70, 95% CI = [-0.73, -0.67]); right panel: t(286) = -11.02, p < 0.001, r = -0.55, 95% CI = [-0.59, -.050]). Shade of blue indicates ten within-observer bins of increasing confidence. Dots represent single observers (N = 32). (b-c) Decoded sensory uncertainty reliably predicts reported confidence. (b) Example observer (S24; left panel: z = -1.67, p = 0.047, ρ= -0.078, 95% CI = [-0.17, 0.014]; right panel: z = -1.52, p = 0.064, ρ = -0.071, 95% CI = [-0.16, 0.021]). Analyses were performed on trial-by-trial data (N = 493); data were binned (ten bins) for visualization only. Error bars represent ± 1 s.e.m. (c) Group average (red line; shaded area represents ± 1 s.e.m.), probability density, and individual correlation coefficients (Left panel: z = -2.17, p = 0.015, ρ = -0.018, 95% CI = [0.035, -0.0018]; right panel: z = -1.53, p = 0.063, ρ = -0.013, 95% CI = [-.030, .0036]). Gray dots indicate observers (N = 32), circle denotes S24.

We next turned to the relationship between confidence and behavioral precision for repeated presentations of the same stimulus. For each observer, we again sorted trials into ten bins of increasing confidence, calculated the mean level of confidence and behavioral variability across all trials in each bin, and computed the partial correlation coefficient between the two (while controlling for stimulus orientation, see Methods). We considered two possible outcomes. If observers account for trial-by-trial fluctuations in internal noise when estimating confidence, as suggested by both the Probabilistic and Response heuristics model, then higher levels of confidence should predict improved behavioral performance. If, on the other hand, observers rely on orientation heuristics to confidence, then we should observe no systematic relationship at all between confidence and behavioral variability. The results revealed that behavior was more precise when confidence was high (Fig. 3a, right; t(286) = -11.02, p < 0.001, r = -0.55, 95% CI = [-0.62, -.046]). This is consistent with both the Probabilistic and Response heuristics model, and argues against an explanation of confidence in terms of orientation heuristics.

To adjudicate between the two remaining hypotheses, we then turned to the brain data. Specifically, we used a probabilistic decoding algorithm^10, 18^ to characterize the degree of uncertainty in perceptual evidence from cortical activity patterns in areas V1-V3. Uncertainty in the cortical stimulus representation (‘decoded uncertainty’) was quantified on a trial-by-trial basis as the width (variance) of a decoded probability distribution (see Methods). Benchmark analyses verified that 1) orientation decoding performance was well above chance levels (Extended Data Fig. 2a, see also ^10^), 2) decoded uncertainty was lower for cardinal compared to oblique stimuli (Extended Data Fig. 2b, see also ^10^), and 3) decoded uncertainty predicted behavioral variability, both within and across stimulus orientations (Extended Data Fig. 2c-d, see also ^10^). Altogether, this confirms that the precision of the observer’s internal sensory evidence was reliably extracted from the patterns of fMRI activity on a trial-by-trial basis.

Do human observers rely on the quality of their internal visual evidence when estimating confidence? To address this question, we computed, for each individual observer, the trial-by-trial rank correlation coefficient between reported confidence and decoded uncertainty (see Fig. 3b for an example observer). The obtained correlation coefficients were subsequently averaged across observers. Per our simulations, we predicted that if confidence is based on sensory uncertainty, then the imprecision in the observer’s sensory evidence, as assessed by the decoder, should predict the confidence judgments of the observer. If, however, confidence is consistent with heuristic computations based on non-sensory sources of noise, then we should observe no relationship between decoded uncertainty and reported confidence at all. Corroborating the Probabilistic model, there was a reliable inverse relationship between decoded uncertainty and behavioral confidence (z = -2.17, p = 0.015, ρ = -0.018, 95% CI = [-0.035, -0.0018]; Fig. 3c, left). To further substantiate this result, we repeated the analysis while controlling for stimulus orientation (see Methods). Although this slightly reduced the strength of the observed relationship between the fidelity of the cortical stimulus representation and reported confidence (z = -1.53, p = 0.063, ρ = -0.013, 95% CI = [-.030, .0036]; Fig. 3c, right), the correlation coefficient reached significance when using smaller numbers of voxels, which respond more strongly to the visual stimulus (Extended Data Fig. 3). Thus, when the cortical representation of a stimulus is more precise, observers consistently report higher levels of confidence, as predicted by the Probabilistic model (and none of the other models). Control analyses verified that these results were robust to variations in the number of voxels selected for analysis (Extended Data Fig. 3), and moreover, could not be explained by eye movement, position or blinks, nor by mean BOLD amplitude (Extended Data Fig. 4). Taken together, these results suggest that human observers rely on a probabilistic representation of the quality of their sensory evidence when judging confidence.

### Sensory uncertainty and subjective confidence in downstream areas

To further test the probabilistic confidence hypothesis, we next asked which downstream regions might read out the uncertainty contained in visual cortical activity so as to compute confidence. Based on our modeling work, we reasoned that if confidence is based on a probabilistic representation of the evidence, then we should be able to find downstream areas whose activity reflects sensory uncertainty, and predicts reported confidence, on a trial-by-trial basis. Specifically, we predicted an inverse relationship in activity between sensory uncertainty and confidence for these regions (cf. Fig. 2d). Thus, under the probabilistic confidence hypothesis, cortical activity should not only increase (decrease) with reduced reliability of the observer’s perceptual evidence, but also decrease (increase) when the observer reports greater levels of decision confidence.

We first focused on those areas that are driven by internal fluctuations in perceptual uncertainty. To identify candidate areas, we performed a whole-brain search. Specifically, we ran a general linear model (GLM) analysis in which we modeled the BOLD signal as a function of the degree of uncertainty decoded from visual cortical representations (in areas V1-V3), while controlling for differences in stimulus orientation (see Methods for further details). We found several clusters downstream of visual cortex where neural activity reliably co-fluctuated with trial-by trial changes in decoded sensory uncertainty (see Fig. 4a for an overview, Supplementary Data for whole-brain maps, and Supplementary Table 1 for a list of clusters). This included the dorsal anterior insula (dAI), dorsal anterior cingulate cortex (dACC), and left rostrolateral prefrontal cortex (rlPFC) – regions that are commonly associated with uncertainty^22^ (dAI), volatility^23, 24^ (dACC) and metacognition^25^ (rlPFC).

**Fig. 4.**
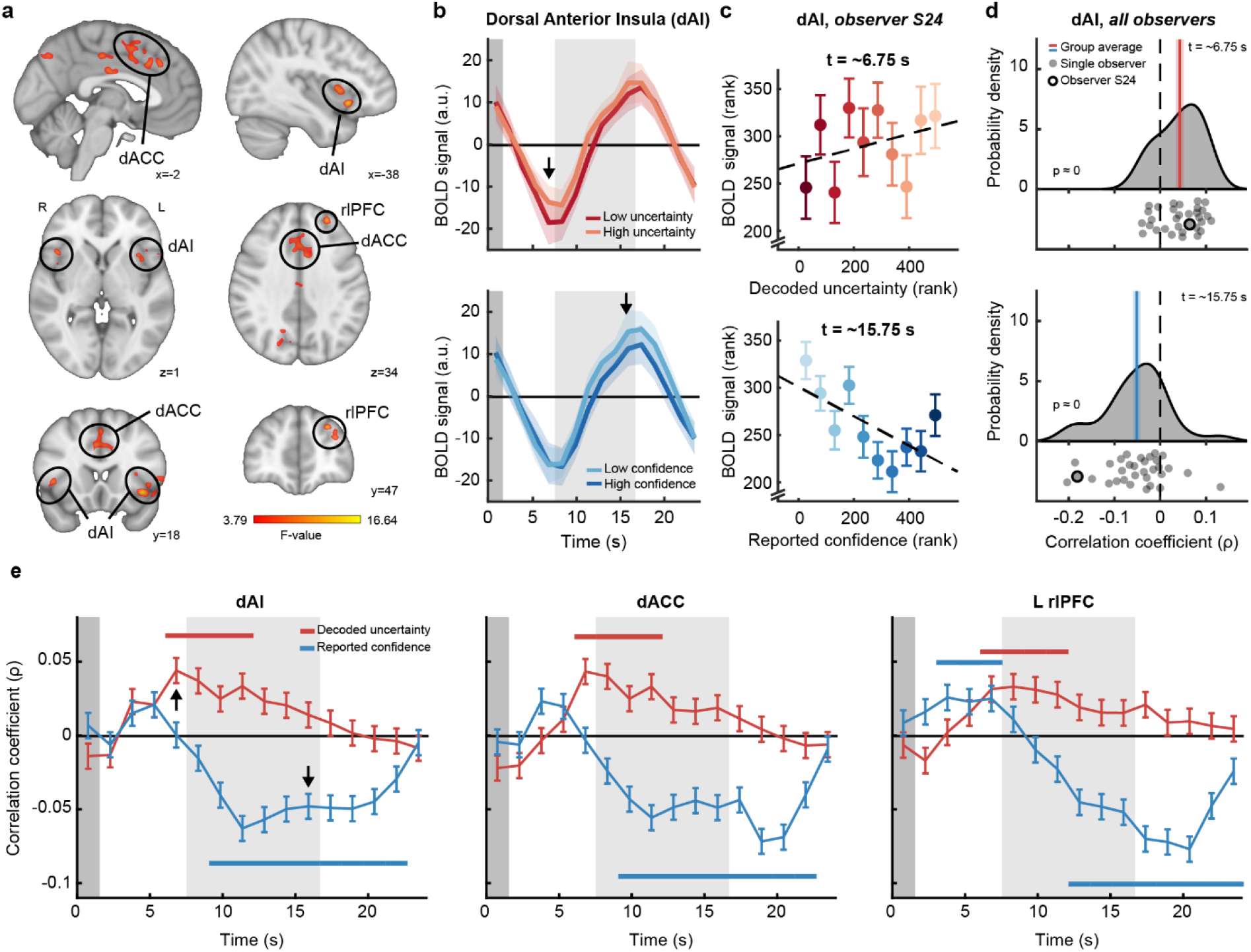
Activity in dorsal anterior insula (dAI), dorsal anterior cingulate cortex (dACC), and left rostrolateral prefrontal cortex (L rlPFC) over time. (a) Downstream clusters significantly modulated by uncertainty decoded from visual cortex (p<0.05 FWER-controlled) (data were masked to exclude occipital cortex; Supplementary Table 1 gives an overview of all activations and see Supplementary Data for whole-brain maps). (b) Cortical response in dAI for high versus low decoded uncertainty (top) and high versus low reported confidence (bottom), averaged across all observers. Trials were binned by a median split per observer. Black arrows indicate the data presented in c and d. (c) Example observer S24. The observer’s cortical response tentatively increases with decoded uncertainty (top panel, permutation test; ρ = 0.065, p = 0.078), and reliably decreases with reported confidence (bottom panel, permutation test; ρ = -0.18, p < 0.001). Trials (N = 493) were binned (ten bins) for visualization only, correlation coefficients were computed from trial-by-trial data. (d) Group average (red/blue line) and correlation coefficients for individual observers (gray dots; N = 32). Shown is the relationship between cortical response amplitude and decoded uncertainty (top) or reported confidence (bottom). Both effects are statistically significant at the group level (permutation test; uncertainty: ρ = 0.043, p < 0.001; confidence: ρ = -0.052, p < 0.001). (e) Group-averaged correlation coefficient between cortical response amplitude and decoded uncertainty (red) or reported confidence (blue). Horizontal lines indicate statistical significance (p<0.05, FWER-controlled). Arrows indicate data in d. (b,e) Dark gray area marks stimulus presentation window, light gray area represents response window. (b-e) Shaded areas and error bars denote ± 1 s.e.m.

We next asked whether these uncertainty-tracking regions would also show a reliable opposite relationship to confidence in their activity, as predicted by the Probabilistic observer model. To address this question, we performed region-of-interest (ROI) analyses within the candidate regions identified by the above whole-brain analysis. First, individual ROIs were created by selecting all uncertainty-driven voxels within predefined anatomical masks corresponding to dAI^26^, dACC^27^, and left rlPFC^28^, using a leave-one-subject-out cross-validation procedure to avoid double dipping^29, 30^ (see Methods for details, and Extended Data Fig. 5 for remaining clusters). For each subject, we then averaged the BOLD signal across all voxels within the ROI. To test whether BOLD activity was reliably modulated by the level of confidence reported by the observer, we performed a GLM analysis (see Methods for model details). This revealed a significant effect of confidence on BOLD activity in all three regions (dAI: F(3,93) = 25.94, p < 0.001, R^2^ = 0.26, 95% CI = [0.12, 0.41]; dACC: F(3,93) = 27.33, p < 0.001, R^2^ = 0.25, 95% CI = [0.10, 0.39]; rlPFC: F(3,90) = 2.88, p = 0.040, R^2^ = 0.01, 95% CI = [-0.03, 0.05]). Thus, it appears that neural activity in dAI, dACC, and rlPFC is affected by both the trial-by-trial imprecision in sensory evidence and the level of confidence reported by the observers.

Having established that activity in dAI, dACC and rlPFC is modulated by confidence, we next investigated the hypothesized inverse relationship between sensory uncertainty and reported confidence on the cortical response in these regions. To illustrate our approach, we first focus on a single ROI (dAI; Fig. 4b). We computed, for each observer, the trial-by-trial correlation coefficient between decoded uncertainty and mean BOLD amplitude within the ROI (after removing the effect of stimulus orientation, see Methods; see Fig. 4c for an example observer), averaged the coefficients across observers (Fig. 4d), and repeated the analysis over time (Fig. 4e, left panel). We also performed the same analysis for reported confidence (Fig. 4b-e), and dACC and rlPFC (Fig. 4e). We discovered, in all three ROIs, a significant positive relationship between decoded uncertainty and cortical activity that was sustained over an extended period of time (Fig. 4e; all p < 0.05, FWER-controlled). Critically, the effect on the cortical response was reversed for confidence (Fig. 4e), and similarly held up over time (all p < 0.05, FWER-controlled). Thus, while the cortical response in dAI, dACC and rlPFC reliably increased with decoded uncertainty, activity in these regions consistently decreased with reported confidence, further corroborating the Bayesian confidence hypothesis. These effects could not be explained by trial-by-trial fluctuations in the participant’s response time (see Extended Data Fig. 6). Especially interesting is that (in dACC and dAI) the positive correlation with uncertainty temporally preceded the negative correlation with reported confidence (dAI: t(31) = -3.05, p = 0.005, d = -0.54, 95% CI = [-0.90, -0.17]; dACC: t(31) = -2.72, p = 0.011, d = -0.39, 95% CI = [-0.75, -0.032]; rlPFC: t(30) = -1.09, p = 0.29, d = -0.20, 95% CI = [-0.57, 0.17]); factoring in the (approximately 4-second) hemodynamic delay inherent in the BOLD response, the effect of sensory uncertainty appears to be roughly time-locked to the presentation (and neural processing) of the stimulus, while the correlation with reported confidence coincides with the time when subjects had to estimate their confidence. These distinct latencies furthermore suggest that decoded uncertainty and reported confidence exert (partially) independent effects on activity in these regions, which was confirmed by a partial correlation analysis (Extended Data Fig. 7). Moreover, cortical activity in all three regions was additionally found to mediate the trial-by-trial relationship between decoded uncertainty and reported confidence (Extended Data Fig. 8), indicating that the effects of sensory uncertainty and subjective confidence on activity in these regions also include a shared component. Taken together, these results are consistent with the Bayesian confidence hypothesis, and suggest that dAI, dACC, and rlPFC are involved in the computation of confidence from a probabilistic representation of the quality of the observer’s sensory evidence.

## Discussion

What computations give rise to the subjective sense of confidence? Here, we tested the Bayesian hypothesis that confidence is computed from a probabilistic representation of information in cortex. We first implemented a Bayesian (Probabilistic) observer model as well as two models using alternative strategies to confidence. This resulted in a set of predictions that we tested using psychophysics and fMRI. Corroborating the Bayesian model, we found that reported confidence reflects behavioral precision, even when stimulus properties such as orientation are held constant. Moreover, probability distributions decoded from population activity in visual cortex predict the level of confidence reported by the participant on a trial-by-trial basis. We furthermore identified three downstream regions, dACC, dAI and rlPFC, where BOLD activity is linked to both the width of the decoded distributions and reported confidence in ways consistent with the Bayesian observer model. Taken together, these findings support recent normative theories, and suggest that probabilistic information guides the computation of one’s sense of confidence.

Earlier work on statistical confidence has manipulated evidence reliability by varying physical properties of the stimulus, such as its contrast. This left open the possibility that observers simply monitor these image features as a proxy for uncertainty^7, 10–12^, without considering an internal belief distribution over the latent variable. For this reason, we held stimulus properties constant, relied on fluctuations in internal noise, and extracted probability distributions directly from cortical activity. Our work shows that the uncertainty decoded from visual activity predicts the level of confidence reported by the observer. No less important, we find that downstream regions commonly associated with volatility in the environment^23, 24^, decision-making^22, 31^, and confidence^25, 32, 33^, represent trial-by-trial fluctuations in both this decoded uncertainty and reported confidence. Altogether, this strongly suggests that not stimulus heuristics, but rather a probabilistic representation of information drives human confidence reports.

While decision confidence is usually studied in the context of binary decisions, we here focused on a continuous estimation task, which requires observers to reproduce a feature of the stimulus. For binary decisions, confidence is normatively defined as a function of the observer’s measurement and decision boundary, in addition to sensory uncertainty, and each of these parameters can vary on a trial-by-trial basis due to internal noise. For the continuous estimation task used here, on the other hand, confidence is more straightforwardly defined as a function of sensory uncertainty, without many additional parameters. This definition makes this task ideally suited for addressing the probabilistic confidence hypothesis. While we specifically focused on uncertainty in continuous estimation, it seems nonetheless likely that the probabilistic nature of the representation will extend to binary choices and other decisions of increasing complexity.

Our findings are also important for understanding how uncertainty is represented in cortex. Previous work has shown that the width of the decoded probability distribution predicts the magnitude of behavioral orientation biases^10^, serial dependence effects in perception^34^, and classification decisions^35^. The current work extends these earlier findings by linking the decoded distributions directly to activity in downstream decision areas and the subjective level of confidence reported by the observer. Taken together, these findings suggest that probability distributions are not only represented in neural population activity, but also used in the brain’s computations.

Earlier work has implicated the involvement of the dACC, dAI, and rlPFC in experimental (objective) manipulations of evidence reliability^23, 33, 36–38^. Our results suggest that these regions similarly track spontaneous (internal) fluctuations in uncertainty, further elucidating their functional role in human decision-making. Thus, it appears that a more general notion of uncertainty is represented in these regions, albeit for different functional purposes. While the representation in dACC may serve to inform internal models and response selection^39–42^, it seems likely that dAI integrates uncertainty with interoceptive and affective information to form a general subjective feeling state^22^. rlPFC, on the other hand, likely plays a key role in the integration of internal uncertainty with contextual information to compute confidence^32, 36, 43–45^.

While we here focused on the link between subjective confidence and the precision of early sensory evidence in visual areas, confidence should additionally reflect uncertainty added by later stages of processing; for instance, when the item is held in visual working memory before observers make their judgment (although the impact of memory imprecision might be relatively small^46^). Indeed, our ideal observer model incorporates these sources of variance in its estimates of confidence (cf. equations 1, 8 and 9, *σ*_*n*_ ^2^). Given the involvement of early visual areas in visual working memory^47, 48^, this predicts that the imprecision in the visual cortical representation during the delay period between stimulus presentation and response should predict the level of confidence reported by the observer, as well. Interestingly, additional analysis of the empirical data revealed that uncertainty decoded from signals during the retention interval indeed reliably predicted subjective levels of confidence (Extended Data Fig. 9). Although our design does not warrant strong conclusions regarding the nature of these signals (see Extended Data Fig. 9 for discussion), these findings are consistent with a model that considers additional sources of variance when judging confidence. It will be interesting for future work to further disentangle the various sources of noise that affect the observer’s decisions and associated levels of confidence.

In conclusion, we showed that behavioral confidence tracks the degree of uncertainty contained in neural population activity in visual cortex, suggesting that human observers have access to and can report about the degree of imprecision in their visual cortical representations of the stimulus. Furthermore, activity in the dACC, dAI and rlPFC is modulated by both this uncertainty and reported confidence in ways predicted by the Bayesian model, suggesting that these regions are involved in the computation of confidence from sensory uncertainty. Taken together, the current results support recent normative theories of confidence and suggest that the subjective feeling of confidence is based on a statistical measure of the quality of one’s evidence.

## Methods

### Participants

32 healthy adult volunteers (age range 19-31, 20 female) with normal or corrected-to-normal vision participated in this study. Sample size (N=32) was based on a power calculation (power = 0.8; α = 0.05). All participants gave informed written consent prior to their participation and received monetary compensation for their participation. The study was approved by the local ethics committee (CMO Arnhem-Nijmegen, the Netherlands). Participants were included based on their ability to perform the task, which was assessed in a separate behavioral training session prior to the experimental sessions.

### Imaging data acquisition

MRI data were acquired on a Siemens 3T MAGNETOM PrismaFit scanner at the Donders Center for Cognitive Neuroimaging, using a 32-channel head coil. For anatomical reference, a high-resolution T1-weighted image was collected at the start of each session (3D MPRAGE, TR: 2300 ms, TI: 1100 ms, TE: 3 ms, flip angle: 8 degrees, FOV: 256 x 256 mm, 192 saggital slices, 1-mm isotropic voxels). B0 field inhomogeneity maps (TR: 653 ms, TE: 4.92 ms, flip angle: 60 degrees, FOV: 256 x 256 mm, 68 transversal slices, 2-mm isotropic voxels, interleaved slice acquisition) were acquired. Functional data were acquired using a multi-band accelerated gradient-echo EPI protocol, in 68 transversal slices covering the whole brain (TR: 1500 ms, TE: 38.60 ms, flip angle: 75 degrees, FOV: 210 x 210 mm, 2-mm isotropic voxels, multiband acceleration factor: 4, interleaved slice acquisition).

### Experimental design and stimuli

Participants performed an orientation estimation task while their cortical activity was measured with fMRI. They completed a total of 22-26 task runs, divided over two scan sessions on separate days. Prior to the experimental sessions, participants extensively practiced the task (2-4 hours) in a separate behavioral session.

Throughout each task run, participants fixated a bull’s eye target (radius: 0.375 degrees) presented at the center of the screen. Each run consisted of 20 trials (16.5 s each), separated by an inter-trial interval of 1.5 s, and started and ended with a fixation period (duration at start: 4.5 s; at end: 15 s). Each trial started with the presentation of the orientation stimulus, which remained on the screen for 1.5 s. This was followed by a 6-s fixation interval, and then two successive 4.5-s response windows (Extended Data Fig. 1). Orientation stimuli were counterphasing sinusoidal gratings (contrast: 10%, spatial frequency: 1 cycle per degree, randomized spatial phase, 2-Hz sinusoidal contrast modulation) presented in an annulus around fixation (inner radius: 1.5 degrees, outer radius: 7.5 degrees, grating contrast decreased linearly to 0 over the inner and outer 0.5 degrees of the radius of the annulus). Stimulus orientations were drawn (pseudo)randomly from a uniform distribution covering the full orientation space (0-179 degrees) to ensure an approximately even sampling of orientations within each run. At the start of the first response window, a black bar (length: 2.8 degrees, width: 0.1 degrees, contrast: 40%) appeared at the center of the screen at an initially random orientation. Subjects reported the orientation of the previously seen grating by rotating this bar, using separate buttons for clockwise and counterclockwise rotation on an MRI-compatible button box. At the start of the second response window, a black bar of increasing width (contrast: 40%, bar width: 0.1-0.5 degrees, linearly increasing) and wrapped around fixation (radius 1.4 degrees) became visible at the center of the screen. Participants indicated their confidence in their orientation judgement by moving a white dot (contrast: 40%, radius: 0.05 degrees) on this continuous confidence scale, using the same buttons for clockwise and counterclockwise as for their orientation response. The mapping of confidence level to scale width (i.e. whether the narrow end of the scale indicated high or low confidence) was counterbalanced across participants. The scale’s orientation and direction (i.e. width increasing in clockwise or counterclockwise direction), as well as the starting position of the dot, were randomized across trials. For both response windows, the bar (scale) disappeared gradually over the last 1 s of the response window to indicate the approaching end of this window. Shortly before trial onset (0.5 s), the fixation bull’s eye briefly turned black (duration: 0.1 s) to indicate the start of the trial. Because we were interested in the effects of sensory uncertainty on cortical activity and confidence, rather than the cortical representation of confidence *per se*, and moreover, reward-related signals might contaminate the representation of sensory information in visual areas^49^, participants received no trial-by-trial feedback about the accuracy of their judgments.

Each scan session also included 1 or 2 functional localizer runs, during which flickering checkerboard stimuli were presented in seven 12-s blocks interleaved with fixation blocks of equal duration. The checkerboard stimuli were presented within the same aperture as the grating stimuli (contrast: 100%, flicker frequency: 10 Hz, check size: 0.5 degrees). Retinotopic maps of the visual cortex were acquired in a separate scan session using standard retinotopic mapping procedures^50–52^.

All visual stimuli were generated on a Macbook Pro computer using Matlab and the Psychophysics Toolbox^53^, and were presented on a rear-projection screen using a luminance-calibrated EIKI LC-XL100 projector (screen resolution: 1024 x 768 pixels, refresh rate: 60 Hz). Participants viewed the screen through a mirror mounted on the head coil.

### Behavioral data analysis

In general, participants finished adjusting their orientation and confidence responses well before the end of the response windows (4.5 s each), taking on average 2761 ± 378 ms (mean ± S.D. across observers) for the orientation response and 2587 ± 313 ms for the confidence response. Trials on which participants did not finish their response by the end of the response window were excluded from further analyses (0-43 out of 440-520 trials). The error in the observer’s behavioral orientation response was computed as the acute-angle difference between the reported and the presented orientation on a given trial. Orientation-dependent shifts (biases) in mean behavioral error were removed by fitting two fourth-degree polynomials to each observer’s behavioral errors as a function of stimulus orientation (see ^10^ for a similar procedure). One polynomial was fit to trials for which the presented stimulus orientation was between 90 and 179 degrees, and the second polynomial was fit to trials on which the presented stimulus orientation was between 0 and 89 degrees. We used the bias-corrected behavioral errors, i.e. the residuals of this fit, in subsequent analyses. Behavioral errors that were more than three standard deviations away from the mean of each participant (after bias correction) were marked as guesses and excluded from further analysis (1-7 out of 440-520 trials). To remove potential session- and subject-specific differences in usage of the confidence scale, confidence ratings were z-scored within sessions.

### Preprocessing of MRI data

The raw functional imaging data were motion-corrected with respect to the middle volume of the middle run of the session, using FSL’s MCFLIRT^54^. The functional data were corrected for distortion using the within-session B0 fieldmap, and aligned to the T1-weighted image obtained during the same scan session. This anatomical (T1-weighted) image was aligned with a subject-specific unbiased template image, created by combining the T1-weighted images from the two sessions, using Freesurfer’s mri_robust_template^55^. Slow drifts in the BOLD signal were removed using FSL’s nonlinear high-pass temporal filter with a sigma of 24 TRs (two trials), corresponding to a cut-off period of approximately 83 seconds.

For all univariate analyses, additional preprocessing steps were performed prior to high-pass filtering. Specifically, non-brain structures were removed using FSL’s BET^56^, and the data were spatially smoothed with a 6-mm Gaussian kernel using FSL’s SUSAN^57^. As the univariate analyses required combining data across subjects, each subject’s anatomical template image was non-linearly registered to MNI152 space using FSL’s FNIRT with a warp resolution of 10 mm isotropic^58^.

A set of nuisance regressors was used to remove residual motion effects and global fluctuations in the BOLD signal. Per session, we defined an intercept regressor per run, 24 motion regressors based on the motion parameters estimated by MCFLIRT (all analyses), and two regressors reflecting the average signal in cerebrospinal fluid (CSF) and white matter (WM) (univariate analyses only). The CSF and WM regressors served to capture global fluctuations in signal intensity and were obtained by first creating WM and CSF masks based on the subject’s anatomical scan data using FSL’s FAST ^59^, and then removing the outer edges from these masks to exclude voxels at the tissue boundaries. For the multivariate and ROI-based univariate analyses, nuisance signals were removed from the BOLD signal prior to further analyses. For the whole-brain univariate analysis, motion, CSF/WM, and intercept regressors were included as covariates in the general linear model (see Whole-brain analysis).

For the multivariate analyses, the ROI (consisting of V1, V2, and V3) was identified on the reconstructed cortical surface. Within this ROI, and in the native space of each participant, we selected for further analysis the 2000 voxels that were activated most strongly by the functional localizer stimulus while surviving a lenient statistical threshold (p < 0.01, uncorrected). Control analyses verified that our results were not strongly affected by the number of voxels selected for analysis (Extended Data Fig. 3). The time series of each selected voxel was subsequently z-normalized with respect to corresponding trial time points in the same run. Activation patterns for each trial were obtained by averaging over the first 3 s of each trial, after adding a 4.5-s temporal shift to account for hemodynamic delay. This relatively short time window was chosen so as to ensure that activity from the behavioral response window was excluded from analysis. For the control analyses of Extended Data Fig. 4, mean BOLD intensity values were calculated by averaging the z-normalized activation values across the selected voxels and time window. The results of Extended Data Fig. 9 were obtained using a sliding window of size 3 s (2 TRs), chosen so as to match the window size of the main analysis. For each window of analysis, the z-normalized activation values were averaged and subsequently fed to the decoding algorithm.

### Multivariate analysis (visual cortex)

#### Decoding algorithm

Trial-by-trial uncertainty in cortical stimulus representations was computed using a generative model-based, probabilistic decoding algorithm^10, 18^, applied to selected voxels in visual cortex (see previous section for voxel selection criteria). The model describes the generative distribution of the voxel activity patterns given a certain stimulus, *p*(**b**|*s*); in other words, the probability that stimulus *s* will evoke activation pattern **b**. The model assumes that, across trials, voxel activity follows a multivariate Normal distribution around the voxel’s tuning curve for orientation. Voxel tuning curves are defined as a linear combination **W*f***(*s*) of 8 bell-shaped basis functions, each centered on a different orientation (cf. ^60^):

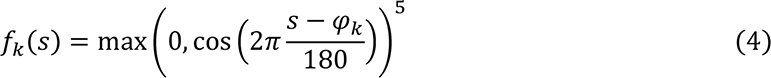

where *s* is the orientation of the presented stimulus and *φ* is the preferred orientation of the *k*-th population. Basis functions were spaced equally across the full orientation space (0-179 degrees) with the first centered at zero degrees. *W*_*iikk*_ is the contribution of the *k*-th basis function to the response of the *i*-th voxel.

The covariance around the voxel tuning curves is described by noise covariance matrix **Ω**:

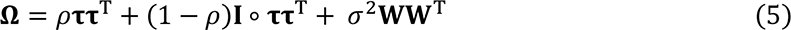

The first term of this covariance matrix describes noise shared globally between all voxels in the ROI, and the second term refers to noise specific to individual voxels (with variance *τ*_*i*_ ^2^ for voxel *i*). The relative contribution of each of these types of noise is reflected in *ρ*. The third term models tuning-dependent noise, i.e. noise, with variance *σ*^2^, shared between voxels with similar orientation preference.

Thus, the generative distribution of voxel responses is given by a multivariate Normal with mean **W*f***(*s*) and covariance **Ω**:

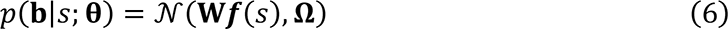

where **θ** = {**W**, **τ**, *ρ*, *σ*} are the model’s parameters. The model’s parameters were estimated in a two-step procedure (see ^10^ for further details). First, the tuning weights **W** were estimated by ordinary least squares regression. In the second step, the noise covariance parameters (*ρ*, *σ*, **τ**) were estimated by numerical maximization of their likelihood.

Model training and testing (‘decoding’) was performed following a leave-one-run-out cross-validation procedure to prevent double-dipping^29^. That is, model parameters were first fit to a training dataset consisting of all but one fMRI run, and the model was then tested on the data from the remaining run. This procedure was repeated until all runs had served as a test set once.

Using the fitted parameters, a posterior distribution over stimulus orientation was computed for each trial in the test set. The posterior distribution is given by Bayes’ rule:

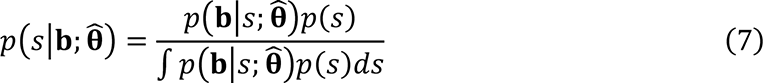

where 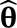 are the estimated model parameters. The stimulus prior *p*(*s*) was flat, given that the stimuli presented in the experiment were uniformly distributed, and the normalizing constant in the denominator was calculated numerically. The circular mean of the posterior distribution was taken as the estimate of the presented orientation on that test trial, and the squared circular standard deviation was used as a measure of the amount of uncertainty in this estimate.

### Statistical procedures

Most of our analyses relied on the computation of a correlation coefficient between two variables. These coefficients were calculated for each individual participant, and then averaged across observers (see below). Based on the assumed relationship between the two variables (linear or monotonic), either Pearson’s or Spearman’s (rank) correlation coefficient was computed. For the analyses using Pearson’s correlation coefficient, the data were visually inspected and appeared to be normally distributed, although this was not formally tested (see Fig. 3a and Extended Data Fig. 4). Decoding accuracy was quantified by computing the circular analog of the Pearson correlation coefficient between the presented and decoded stimulus orientation. To test for an oblique effect in decoded uncertainty, we first calculated for each presented stimulus orientation its distance to the nearest cardinal (i.e. horizontal or vertical) orientation, and then computed the Spearman correlation coefficient between this measure and decoded uncertainty. To test the relationship between reported confidence and decoded uncertainty (independent of stimulus orientation), we first removed orientation-dependent shifts in decoded uncertainty and confidence by modeling confidence (decoded uncertainty) as a quadratic (linear) function of distance to cardinal (see Extended Data Fig. 10); the rank correlation coefficient between confidence and decoded uncertainty was subsequently computed on the residuals of these fitted functions. After obtaining correlation coefficients for each individual observer *i*, the coefficients were Fisher transformed and a weighted average was computed across observers. Specifically, the weight of the *i*-th correlation coefficient was calculated as *w*_*i*_ = 1/*ν*_*i*_, where *ν*_*i*_ is the variance of the Fisher transformed correlation coefficient^61^. For the Pearson correlation, *ν*_*i*_ is given by 1/(*n*_*i*_ – 3) (where *n*_*i*_ is the number of trials), and for the Spearman correlation *ν*_*i*_ = 1.06/(*n*_*i*_ – 3)^62^. Weights were adjusted for the additional degrees of freedom lost due to stepwise correction for the oblique effect in decoded uncertainty or reported confidence by subtracting 1 (for linear correction) or 2 (for quadratic correction) from the denominator in the variance term. The significance of the coefficients was assessed using a Z-test, testing specifically for effects in the direction predicted by the ideal observer models. The average of the Z-transformed values was translated back to the correlation scale for reporting purposes. Similar procedures were used for the control analyses of Extended Data Fig. 4. For these control analyses, we additionally performed equivalence tests (using the two one-sided tests procedure^63^), comparing against a smallest effect size of interest of ρ = 0.1 (see ^64^ for further rationale).

Some of our analyses required the computation of a dispersion measure (i.e., behavioral variability). For these analyses, each participant’s data were first divided into ten equal-size bins, based on either reported level of confidence or decoded uncertainty, and summary statistics were computed across all trials in a given bin. Behavioral variability was computed as the squared circular standard deviation of (bias-corrected) estimation errors across all trials in each bin, and the average level of confidence or decoded uncertainty was quantified by computing the statistical mean. To test the relationship between behavioral variability and confidence (or decoded uncertainty), we used a multiple linear regression analysis. Independent variables were level of confidence (or decoded uncertainty), and the absolute distance between the stimulus and the nearest cardinal axis (mean across trials in each bin). We also included subject-specific intercepts. The dependent variable was behavioral variability. Partial correlation coefficients were computed from the binned data and significance was assessed using t-tests, testing for effects in the direction predicted by the ideal observer models. Control analyses verified that our results did not strongly depend on the number of used bins, nor on the specific shape of the function used to model the effect of stimulus orientation on confidence (or decoded uncertainty).

### Univariate analyses (whole-brain and ROI-based)

#### Whole-brain analysis

To identify brain regions that are modulated by sensory uncertainty, we used a whole-brain general linear model (GLM) approach. A GLM can be written as ***y*** = **X*β*** + ***ε***, where ***y*** represents the timeseries of a single voxel, **X** is referred to as the design matrix (or model), ***β*** is a vector of model parameters, and ***ε*** represents the residuals.

We constructed a model of task-related activity based on three components: 1) a 1.5-s boxcar function time-locked to the stimulus onsets of all excluded trials, with height one, 2) a 1.5-s boxcar function time-locked to the stimulus onsets of all included trials, with height one, 3) a 1.5-s boxcar function time-locked to the stimulus onsets of all included trials, with its height equal to the decoded uncertainty on that trial (linearly corrected for trial-by-trial differences in stimulus orientation, cf. Multivariate analysis). Each boxcar function was convolved with a canonical hemodynamic response function (HRF) and temporal and dispersion derivatives of the HRF (SPM’s informed basis set), yielding a total of nine regressors to include in the design matrix. The derivatives were added for additional model flexibility regarding the shape and latency of the BOLD response. In addition to the task-related regressors, we further included nuisance regressors (24 motion regressors and 2 CSF/WM regressors per session) and run-specific intercepts (see Preprocessing of fMRI data) to improve overall model fit.

The model **X** was fit to each subject’s timeseries, separately for each voxel, to obtain a set of parameter estimates ***β̂*** Subject-level analyses were performed using SPM12, because of its increased efficiency (relative to FSL) when performing the GLM analysis on concatenated data rather than individual runs. The resulting subject-level ***β̂*** maps were then transformed from subject-specific to standard space (MNI152) to allow for comparison and combination of estimates across subjects. We were specifically interested in the effect of decoded uncertainty on the BOLD response, which was modeled by the three regressors corresponding to the third boxcar function. The combined explanatory power of the three regressors was quantified by computing an F-statistic over the corresponding ***β*** estimates (across subjects). To calculate p-values, a sign-flip test (5000 permutations) was performed in combination with threshold-free cluster enhancement (TFCE)^65^, using FSL’s randomise^66^. The family-wise error rate (FWER) was controlled by comparing the true voxel-wise TFCE scores against the null distribution of the maximum TFCE score across voxels^65, 67^.

### ROI analysis

Brain regions modulated by perceptual uncertainty were selected and further investigated as follows. ROIs were defined using existing anatomical atlases, combined with a functional parcellation based on the whole-brain GLM analysis (as described in more detail above). Specifically, within a given (anatomical) ROI, we selected voxels modulated by decoded uncertainty using the GLM analysis, while applying a leave-one-subject-out procedure^30^ to avoid double-dipping^29^. This led to the definition of eight ROIs, for each participant individually: 1) dorsal anterior insula (using the functional parcellation by Chang et al.^26^, mirrored to obtain bilateral labels, retrieved from Neurovault: https://identifiers.org/neurovault.collection:13), 2) left rostrolateral prefrontal cortex (frontal pole label, Harvard-Oxford cortical atlas^28^, trimmed to include the left hemisphere only), 3) dorsal anterior cingulate cortex (bilateral RCZa and RCZp labels, Neubert cingulate orbitofrontal connectivity-based parcellation^27^), 4) precuneus (precuneus label, Harvard-Oxford cortical atlas^28^), 5) supplementary motor area (SMA label, Sallet dorsal frontal connectivity-based parcellation^68^, mirrored to obtain bilateral labels), 6) dorsal perigenual anterior cingulate cortex (bilateral area 32d, Neubert cingulate orbitofrontal connectivity-based parcellation^27^), 7) ventral posterior cingulate cortex (bilateral area 23ab labels, Neubert cingulate orbitofrontal connectivity-based parcellation^27^), 8) dorsal posterior cingulate cortex (bilateral CCZ labels, Neubert cingulate orbitofrontal connectivity-based parcellation^27^). For ROIs 2 and 8, some of the leave-one-out, GLM-based masks did not contain any voxels. The corresponding data were excluded from further analyses for the respective ROIs (for ROI 2: 1 subject, ROI 8: 2 subjects). The BOLD signal was averaged over all voxels within a given ROI.

Having defined our ROIs, we then proceeded to investigate the effects of confidence in these regions. We did this in two different analyses. To assess the degree to which confidence modulated the BOLD response in each ROIs, we performed a GLM analysis. The model structure was similar to the whole-brain univariate analysis, including three 1.5-s boxcar functions time-locked to stimulus onset: one for excluded trials (height one), one for included trials (height one), and one to model the effect of confidence (included trials only; height equal to confidence value on that trial, quadratically corrected for trial-by-trial differences in stimulus orientation, cf. Multivariate analysis). These boxcar functions were each convolved with SPM’s informed basis set (canonical HRF and its temporal and dispersion derivatives), and nuisance regressors (24 motion and 2 CSF/WM regressors per session). Run intercepts were also added.

To further investigate the magnitude and directionality of effects of reported confidence and decoded uncertainty over the course of a trial (without a priori assumptions regarding the shape or timing of the BOLD response), we also performed a trial-by-trial correlation analysis. Specifically, we computed the Spearman correlation coefficient between BOLD intensity and decoded uncertainty or reported confidence for each TR in the trial. Orientation-dependent changes in decoded uncertainty and confidence were first removed by modeling confidence (decoded uncertainty) as a quadratic (linear) function of distance to cardinal (cf. Multivariate analysis), and the correlation coefficient was computed using the residuals of this fit. For the control analyses presented in Extended Data Fig. 6, response time effects in the BOLD signal were removed by modeling BOLD intensity at each timepoint (relative to stimulus onset) as a linear function of the time it took for the observer to 1) respond to the presented orientation and 2) report confidence on that trial, and the correlation coefficient between BOLD intensity and decoded uncertainty or confidence was computed on the residuals of this fit. For the partial correlation analyses reported in Extended Data Fig. 7, both reported confidence (or decoded uncertainty) and BOLD intensity at each timepoint were modeled as a linear function of uncertainty (or confidence). The residuals of these fits were used to compute the (Spearman) correlation coefficient between confidence (or uncertainty) and the BOLD signal for each timepoint. The single-subject correlation coefficients were Fisher transformed, and a weighted average was computed across observers (cf. Multivariate analysis). Statistical significance was assessed using two-tailed permutation tests, in which uncertainty (or confidence) values were permuted across trials (1000 permutations). To control for multiple comparisons (FWER) we compared against the null distribution of the maximum correlation coefficient across timepoints (cf. Whole-brain analysis). Finally, we tested whether there was a significant difference in latency between the effects of confidence and uncertainty on the BOLD signal in each ROI. To this end we determined, for each subject individually, the (within-trial) timepoint at which the correlation coefficient between BOLD and uncertainty (confidence) was most strongly positive (negative). We then performed a paired t-test on these values, comparing between uncertainty and confidence.

To investigate whether activity in downstream areas mediates the relationship between decoded uncertainty and reported confidence, we performed the following analysis. We first modeled both confidence and uncertainty as a linear function of the BOLD signal in a given ROI and at a given (within-trial) timepoint. We then took the residuals of these model fits and computed the Spearman correlation coefficient between the (residual) uncertainty and confidence values. If the selected ROIs mediate the relationship between uncertainty and confidence, the residual correlation coefficient should be smaller in magnitude than the baseline correlation coefficient between uncertainty and confidence (i.e., not controlled for activity in downstream areas; as reported in Fig. 3c). To quantify the mediating effect of the BOLD signal in each ROI at each timepoint, the single-subject correlation coefficients were Fisher transformed, and we subtracted from these values the (Fisher transformed) baseline correlation coefficient between uncertainty and confidence. Finally, a weighted average was computed across observers (cf. Multivariate analyses). Statistical significance of the predicted reduction in correlation strength was assessed using a permutation test, in which BOLD values were permuted across all trials (following otherwise similar procedures as for the whole-brain and main ROI-based analyses).

### Eyetracking data

Eye tracking data were acquired using an SR Research Eyelink 1000 system for 62 out of 64 sessions. For 11 of these sessions, data were collected for 4-12 runs (out of a total of 10-13) due to technical difficulties with the eye-tracking system. Gaze position was sampled at 1 kHz. Blinks and saccades were identified using the Eyelink software and removed. Eye fixations shorter than 100 ms in duration were similarly identified and removed. Any blinks of duration >1000 ms were considered to be artifacts and removed. For some trials, the quality of eye tracking data was of insufficient quality (as indicated by a high proportion of missing data points). This was identified by computing the percentage of missing (gaze) data points in a time window starting 4.5 seconds before stimulus onset and ending 4.5 seconds after stimulus offset, but excluding the stimulus window itself. Trials were excluded from further analysis if the percentage of missing data points within this pre- and post-stimulus window exceeded 50%. Based on this criterion, 3.84 % ± 1.69 % (mean ± S.D.) of trials were excluded from further analysis. Data were band-pass filtered using upper and lower period cutoffs of 36 s and 100 ms, respectively. The median gaze position per run was computed and subtracted from all data points within that run. All measures of interest were computed during stimulus presentation only, i.e. over the first 1.5 seconds of each trial. Mean eye position was obtained by first computing the mean x- and y-coordinate of the gaze data, and then taking the absolute distance from this position to the central fixation target. The proportion of blinks was computed as the fraction of time labeled as blinks; this included saccades immediately preceding or following a blink. A break from fixation occurred when the absolute distance between gaze position and the central fixation target was more than 1.5 degrees of visual angle. The proportion of fixation breaks was computed as the fraction of time labeled as such.

### Ideal observer models

#### Model description

We implemented three different observer models, which make identical decisions but differ in how they compute confidence from internal signals. Model 1 takes a statistical approach and computes confidence from the degree of imprecision in the orientation judgment. Models 2 and 3 use heuristic strategies; model 2 uses features of the stimulus as a cue to confidence, and model 3 bases confidence on the magnitude of the observed error in the response. We call these the Probabilistic (Bayesian), Stimulus heuristics and Response heuristics observer, respectively (see also Fig. 1).

The observer’s task is to infer the stimulus from incoming sensory signals. These signals are noisy, so that there is no one-to-one mapping between a given stimulus *s* and its measurement *m*. Rather, the relationship between stimulus and measurements is described by a probability distribution *p*(*m*|*s*). We assume that across trials, the sensory measurements follow a (circular) Gaussian distribution centered on the true stimulus *s*, with variance *σ*_*m*_^2^(*s*):

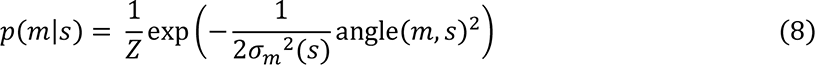

where *Z* is a normalization constant.

We make a distinction between three sources of measurement noise: stimulus-dependent sensory noise 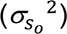, stimulus-independent sensory noise 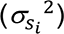, and non-sensory (downstream) noise (*σ*_*n*_ ^2^). The total amount of measurement noise equals the sum of the three noise components:

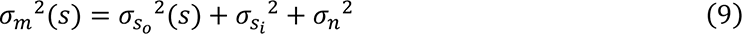

The stimulus-dependent component 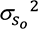 represents Gaussian noise that varies in magnitude as a function of stimulus orientation. Specifically, human behavioral orientation judgments tend to be more precise for cardinal than oblique orientations^19, 69^, and we model this oblique effect in orientation perception as a rectified sine function^20^:

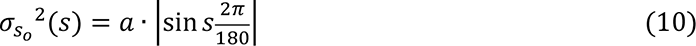

where *a* is the amplitude of the oblique effect. The stimulus-independent component *σ*_*si*_ ^2^ models any remaining sources of (Gaussian) sensory noise. Its magnitude varies randomly over trials, and we model the across-trial distribution of *σ*_*si*_ ^2^ as a gamma distribution:

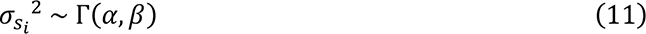

where *α* represents the shape parameter and *β* represents the rate parameter. Finally, the non-sensory (downstream) noise component, *σ*_*n*_ ^2^, is of constant magnitude and captures (Gaussian) noise that arises beyond the initial stages of sensory processing, but prior to the decision, for example when the item is held in working memory or in processing steps downstream of sensory areas V1-V3.

To infer which stimulus likely caused their sensory measurement, the observers use full knowledge of the generative model. Specifically, each observer inverts the generative model using Bayes’ rule. Assuming a flat stimulus prior, the posterior distribution is proportional to the likelihood function:

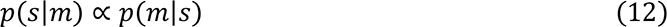

All three observer models take the mean of the posterior distribution as their internal sensory estimate *ŝ* of the presented stimulus. This is the optimal solution for a squared-error loss function^70^. The observer’s internal estimate of orientation is subsequently translated into an overt behavioral (motor) response *r*. The transformation from internal estimate into a motor response is noisy. Thus, the behavioral response *r* for the observer models is given by:

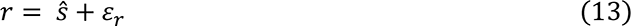

where *ε*_*r*_ is a zero-mean (circular) Gaussian noise variable with variance *σ*_*r*_ ^2^.

The three model observers differ in how they compute confidence. The Bayesian or Probabilistic observer computes confidence as a function of the expected response error. Specifically, this observer assumes a (circular) squared-error loss function and computes confidence as the inverse of the expected loss (Equation 1):

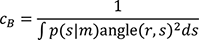

Replacing this direct mapping with any other monotonically decreasing function does not qualitatively change any of the predictions for this model. Thus, for the Bayesian observer, confidence is based (in part) on the posterior probability distribution over the stimulus.

The Stimulus heuristics observer uses the estimated orientation of the stimulus as a cue to uncertainty and confidence. That is, this observer knows that behavior tends to be more precise for cardinal than oblique orientations, and simply exploits this knowledge in their confidence judgments (Equation 2):

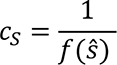

where the function *f*(*s*) takes the shape of the oblique effect (cf. Equation 10):

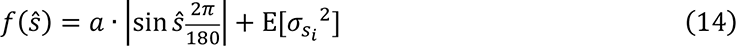

The Response heuristics observer bases confidence on the observed error in the motor response. Specifically, the observer simply notices the difference between the overt response *r* and internal estimate *ŝ*, and adjusts confidence accordingly. We quantified confidence for this model observer as the inverse of the squared acute-angle distance between the internal orientation estimate and the external response (Equation 3):

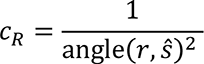

### Simulations

We simulated 50,000 trials for each of the three model observers. Stimulus orientations were drawn from a uniform distribution on the interval [0-179°]. Sensory measurements were randomly sampled from the generative model as described above (Equations 8-11), with *a* = 20, *σ*_*d*_ ^2^ = 5, *α* = 10, and *β* = 1. The normalization constant *Z* was computed numerically. Probabilistic inference proceeded with full knowledge of the parameter values and according to Equation 12. Behavioral responses were obtained using Equation 13 and with *σ*_*r*_ ^2^ = 5. Confidence judgments were obtained using Equations 1-3 and 14. To obtain a reasonable range of confidence values, a constant (of value 1) was added to the denominator of Equation 3. Confidence ratings were z-scored per observer to ensure that they would all fall on the same scale. Sensory uncertainty was quantified as:

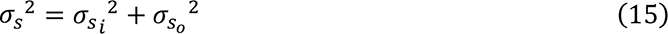

Data were preprocessed following the procedures described in Behavioral data analysis. Similar to the empirical analyses, orientation-dependent shifts in confidence judgments, behavioral variability or sensory uncertainty were removed. For data visualization, simulated data were divided over 10 equal-sized bins of increasing confidence (Fig. 2a-b) or sensory uncertainty (Fig. 2c-d), and the mean confidence level, variance of behavioral errors (Fig. 2a-b), and mean level of sensory uncertainty (Fig. 2c-d) were computed across all trials in each bin.

## Data availability statement

The data are available from the corresponding author upon request.

## Code availability statement

All custom code is available from the corresponding author upon request. Custom code for the probabilistic decoding technique can also be found at https://github.com/jeheelab/TAFKAP

## Acknowledgements

We thank A. Sanfey and R. Cools for helpful discussions, C. Beckmann for advice on statistical analyses, and P. Gaalman for MRI support. This work was supported by European Research Council Starting Grant 677601 (to J.F.M.J.). The funder had no role in study design, data collection and analysis, decision to publish or preparation of the manuscript.

## Author contributions

L.S.G., R.S.v.B. and J.F.M.J. conceived and designed the experiments. L.S.G. collected data. L.S.G. analyzed data, with help from J.F.M.J. L.S.G. and J.R.H.C. constructed ideal observer models, with help from J.F.M.J. L.S.G., J.R.H.C., R.S.v.B. and J.F.M.J. wrote the manuscript.

## Citation diversity statement

We quantified the gender balance of works cited in the main text of this paper (n = 44, excluding self-citations) by manual gender classification of the first and last authors. Among the cited works there are 4.5% single-author male, 72.7% male-male, 2.3% male-female, 15.9% female-male, and 4.5% female-female publications. Expected proportions computed from publications in five top neuroscience journals (as reported in ^71^) are 55.3% male-male, 10.2% male-female, 26.2% female-male, and 8.3% female-female.

## Extended data figures

**Extended Data Fig. 1.**
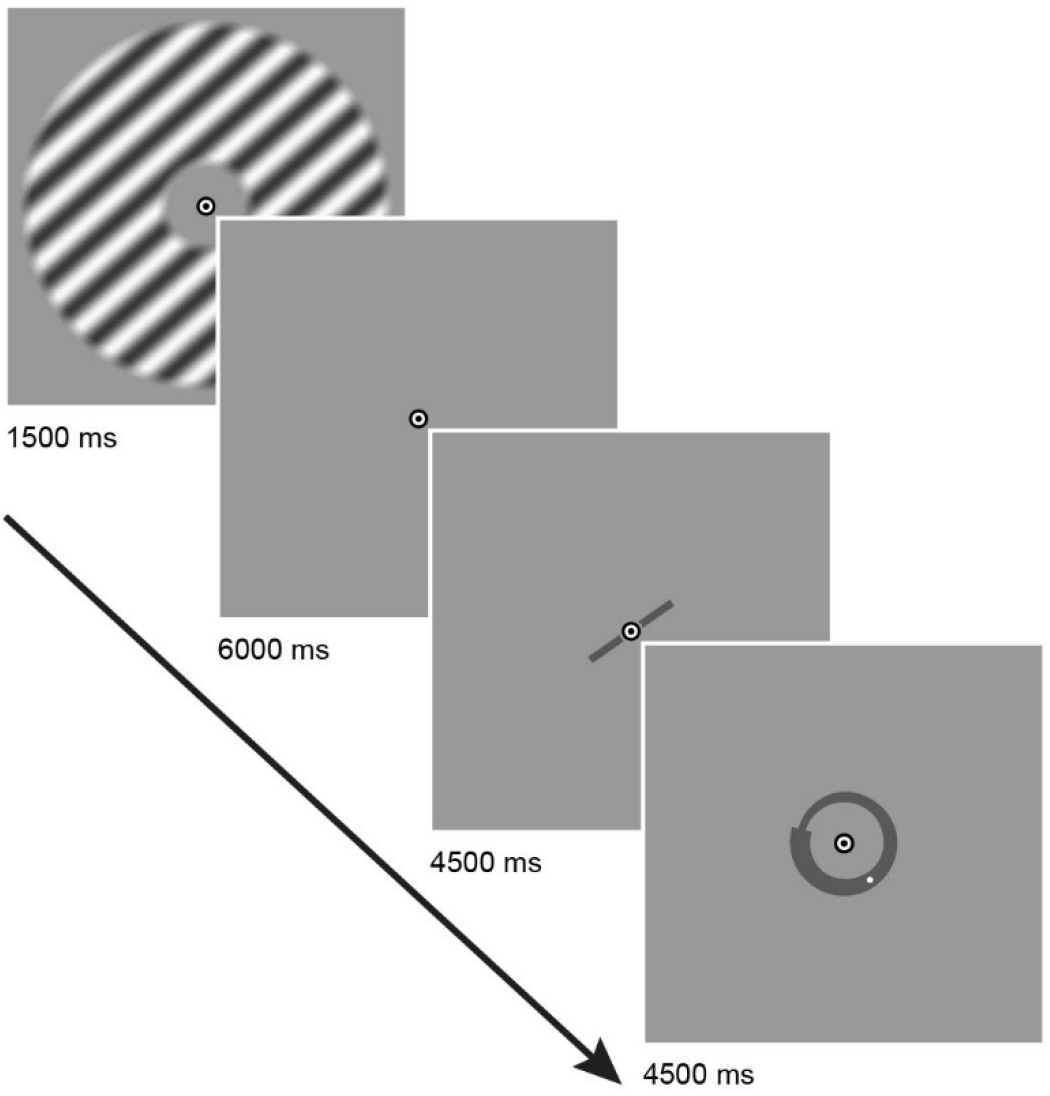
Trial structure. Each trial started with the presentation of an oriented grating (1500 ms) followed by a 6000-ms fixation interval and two 4500-ms response intervals, during which the participant first reported the orientation of the previously seen stimulus by rotating a bar, and then indicated their level of confidence in this judgment on a continuous scale. Trials were separated by a 1500-ms intertrial interval. Stimulus, response bar and confidence scale are not drawn to their true scale and contrast.

**Extended Data Fig. 2.**
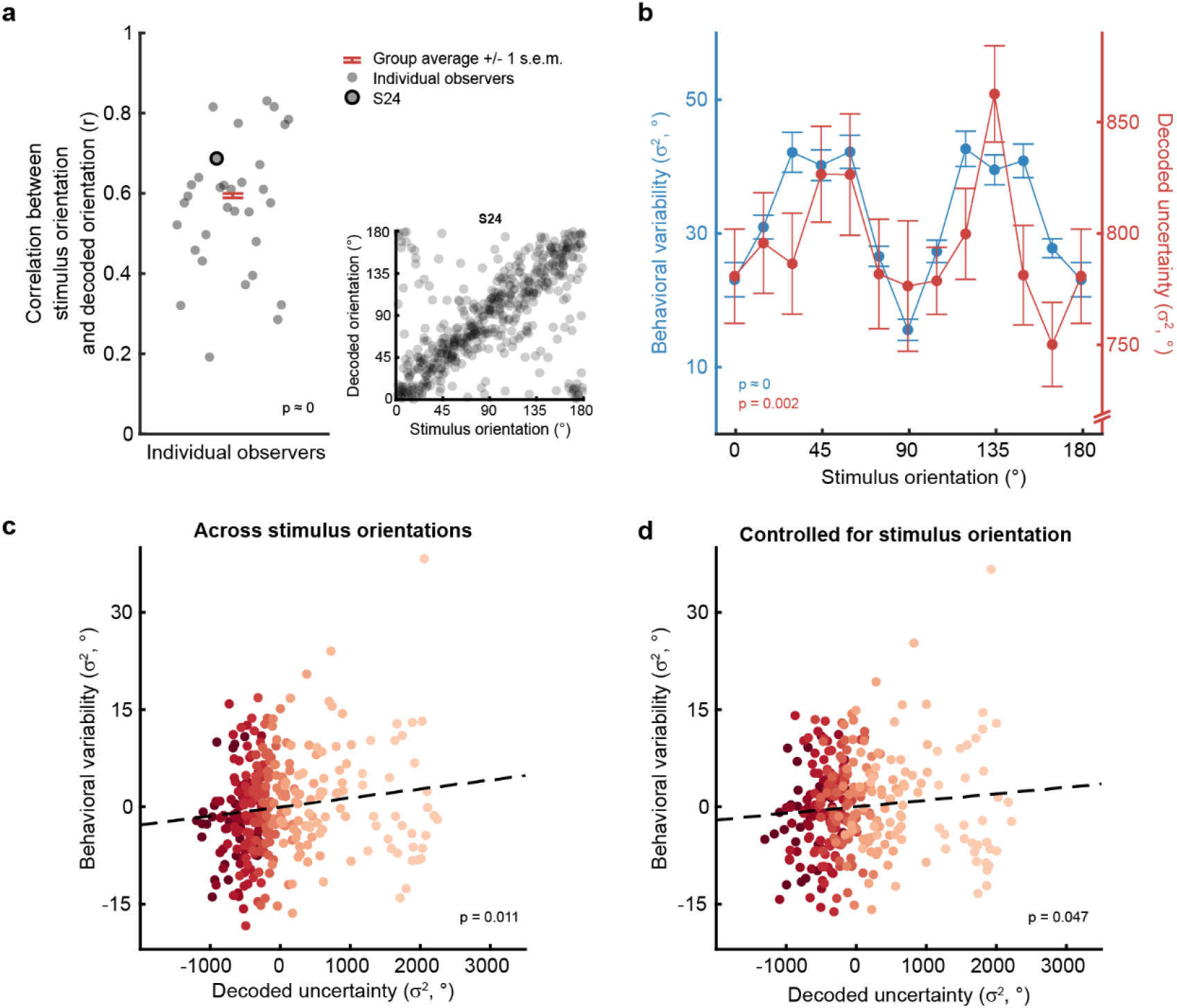
Orientation and uncertainty decoding performance. The orientation of the presented stimulus, and associated uncertainty, decoded from activity patterns in areas V1-V3. (a) Orientation decoding performance was quantified by means of the circular equivalent of the Pearson correlation coefficient between presented and decoded orientations. Correlation coefficients were computed for each subject individually and then averaged across subjects (N = 32). Presented and decoded orientations were significantly correlated (z = 83.58, p < 0.001, r = 0.60, 95% CI = [0.58, 0.61]). (b-d) To assess the degree to which the decoder captured uncertainty contained in neural population activity, we compared decoded uncertainty to behavioral variability, the rationale being that a more precise representation in cortex should also result in more precise behavioral estimates (see also ^10^). (b) Corroborating our approach, we found that decoded uncertainty was greater for oblique compared to cardinal orientation stimuli (correlation distance-to-cardinal and decoded uncertainty: z = 2.95, p = 0.002, ρ = 0.025, 95% CI = [0.0083 0.041]). This finding was paralleled by the imprecision in observer behavior (correlation distance-to-cardinal and behavioral variability: t(287) = 13.60, p < 0.001, r = 0.63, 95% CI = [0.55, 0.69]). (c-d) In addition, behavioral orientation responses were more precise when the decoded probability distributions indicated greater certainty in cortex, (c) both across orientation stimuli (correlation decoded uncertainty and behavioral variability: t(287) = 2.30, p = 0.011, r = 0.13, 95% CI = [0.019, 0.25]), and (d) when controlling for orientation (t(286) = 1.68, p = 0.047, r = 0.099, 95% CI = [-0.017, 0.21]). Altogether, this further underscores the validity of the decoding approach and shows that decoded uncertainty reliably characterizes the degree of imprecision in cortical representations of the stimulus (see ^10, 18^ for further proof of this approach). Note that these are partial residual plots, which is why the data is centered around 0. Error bars (a-b) represent ± 1 s.e.m. (c-d) Shades of red indicate ten equal-size bins of increasing decoded uncertainty, dots represent individual observers (N = 32).

**Extended Data Fig. 3.**
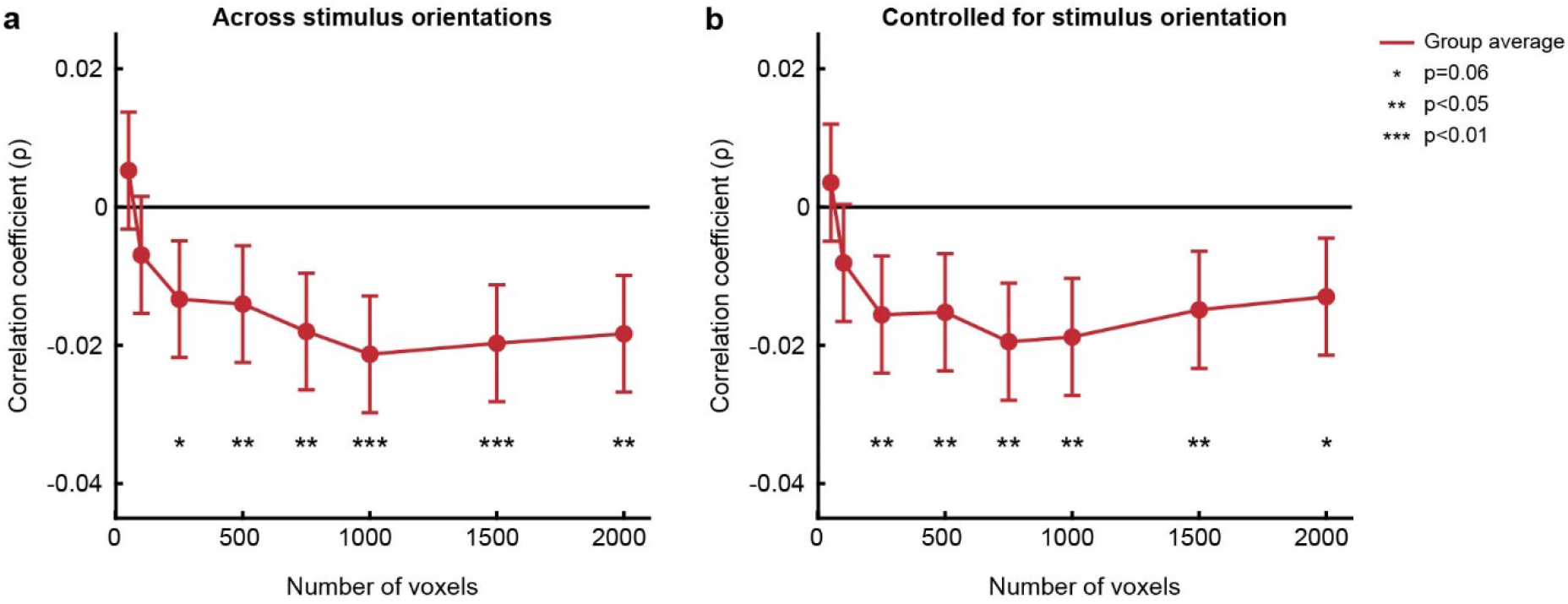
Relationship between decoded uncertainty and reported confidence across different numbers of voxels. Correlation coefficients between decoded uncertainty and reported confidence as a function of the number of voxels included in the ROI, both across all orientations (a) and after removing the effect of stimulus orientation (b). Voxels within V1-V3 were ranked and selected for multivariate analysis based on their response to the visual localizer stimulus (see Methods), using a lenient statistical threshold of p<0.01, uncorrected. The results proved reasonably robust to variations in the number of voxels selected for analysis. Dark red line indicates group average correlation coefficients, error bars denote ± 1 s.e.m.

**Extended Data Fig. 4.**
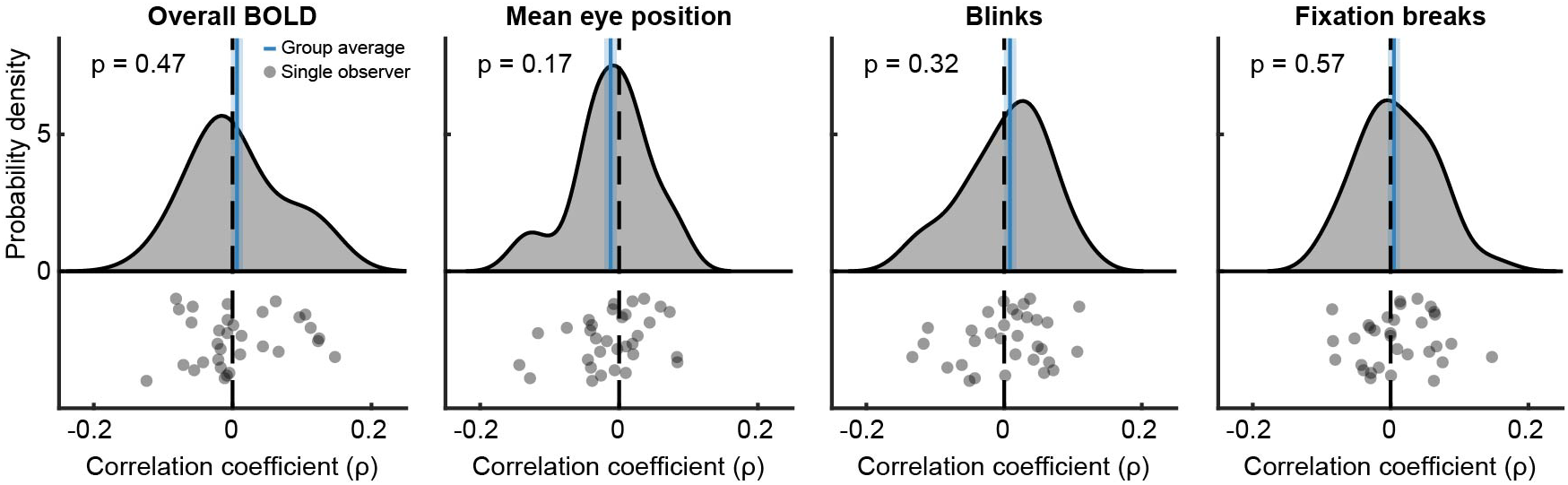
No effects of overall BOLD or eyetracking measures on confidence. Reported confidence is not significantly correlated with the mean BOLD response to the stimulus in areas V1-V3 (z = 0.73, p = 0.47, ρ = 0.0062, 95% CI = [-0.010, 0.023]; equivalence test: z = -0.094, p < 0.001), nor with mean eye position (mean absolute distance to screen center; z = -1.38, p = 0.17, ρ = -0.012, 95% CI = [-0.030, 0.0051]; equivalence test: z = -0.088, p < 0.001), eye blinks (z = 0.99, p = 0.32, ρ = 0.0087, 95% CI = [-0.0086, 0.026]; equivalence test: z = -0.11, p <0.001), or the number of breaks from fixation during stimulus presentation (z = 0.57, p = 0.57, ρ = 0.0050, 95% CI = [-0.012 0.022]; equivalence test: z= -0.11, p < 0.001), suggesting that participants did not rely on heuristics in terms of eye position (‘did I look at the stimulus?’) or eye blinks (‘were my eyes closed?’) for reporting confidence. It furthermore rules out simple heuristic explanations in terms of attentional effort (‘my mind was elsewhere’, ‘I didn’t really try that hard’), as the mean BOLD response to the stimulus tends to increase with attention in these areas^72^. Shaded blue represents ± 1 s.e.m. Gray dots denote individual observers (N = 32).

**Extended Data Fig. 5.**
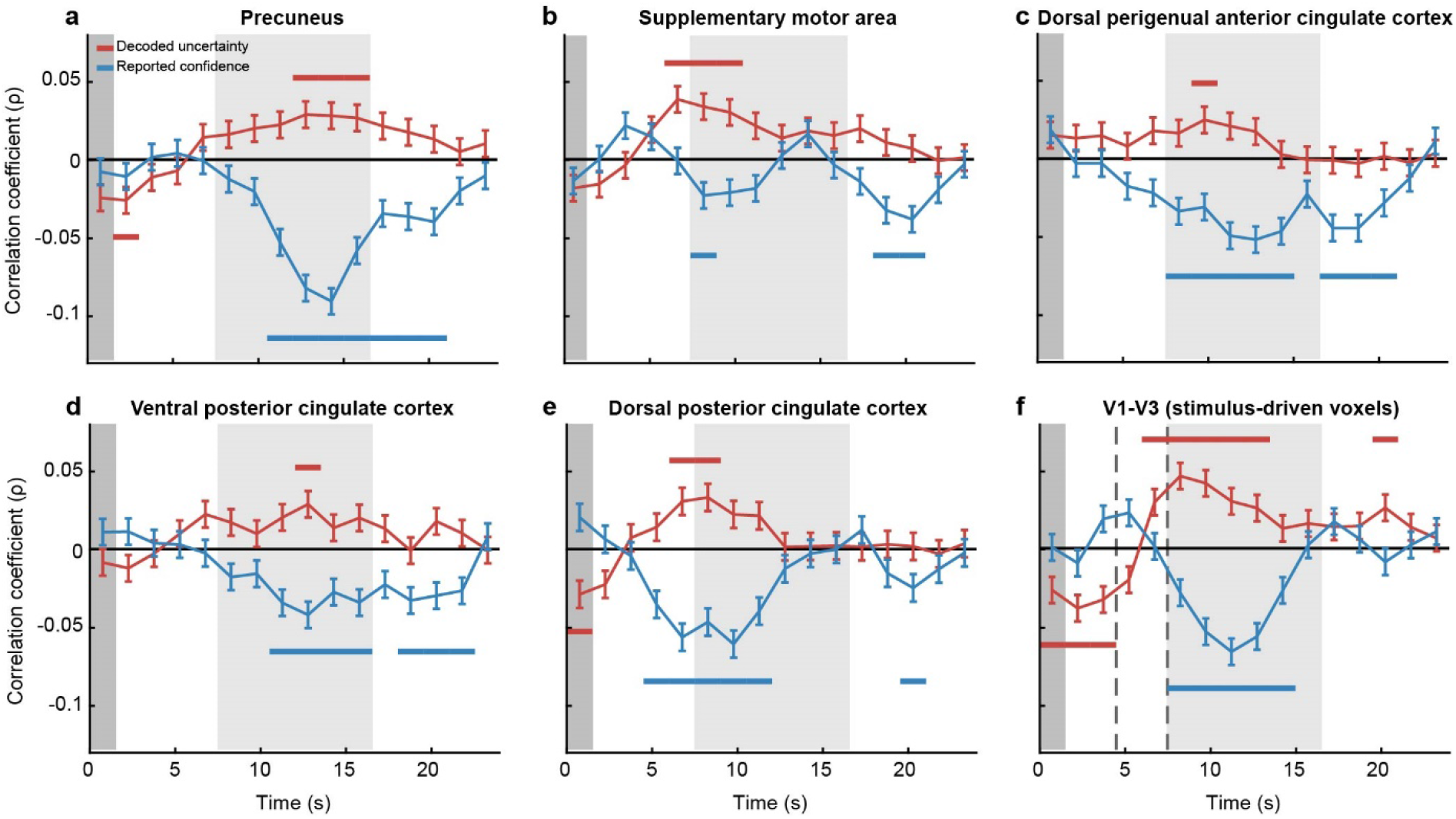
Effects of decoded uncertainty and reported confidence on the BOLD response in precuneus, supplementary motor area, dorsal perigenual anterior cingulate cortex, ventral posterior cingulate cortex, dorsal posterior cingulate cortex, and stimulus-driven voxels in V1-V3. Group-average correlation coefficients for the relationship between decoded uncertainty and BOLD contrast, and reported confidence and BOLD contrast, in six ROIs. (a) In precuneus, the effects of both decoded sensory uncertainty and reported confidence on BOLD peaked around the same time, i.e. during the second half of the response window. This finding is consistent with previous work suggesting that precuneus may represent uncertainty in memory but not in perception^73–75^. (b) In supplementary motor area, both decoded uncertainty and reported confidence modulated cortical activity relatively early in the response window, while the effects of confidence lingered until after observers gave their response. (c-d) In dorsal perigenual anterior cingulate cortex and ventral posterior cingulate cortex, decoded uncertainty had a moderate effect on the BOLD response. Reported confidence modulated cortical activity during as well as shortly after the response window. (e) In dorsal posterior cingulate cortex, the modulatory effect of both decoded uncertainty and reported confidence on the cortical response was largest around the onset of the response window. (f) Stimulus-driven voxels in early visual cortex were modulated by both decoded uncertainty and reported confidence, most notably during the first portion of the response interval. Given the timing of the effect (and taking into account the hemodynamic delay), this likely does not reflect uncertainty in the sensory representation *per se*, but is consistent with anticipatory processes or working memory-related signals potentially influenced by the imprecision in the cortical stimulus representation^76–78^. Please note there is no net effect of uncertainty on the overall (univariate) BOLD response during the decoding window (stimulus presentation; dashed lines). (a-f) Horizontal lines indicate statistical significance (p<0.05, FWER-controlled). Error bars represent ± 1 s.e.m. Dark gray area marks stimulus presentation window, light gray area marks response window.

**Extended Data Fig. 6.**
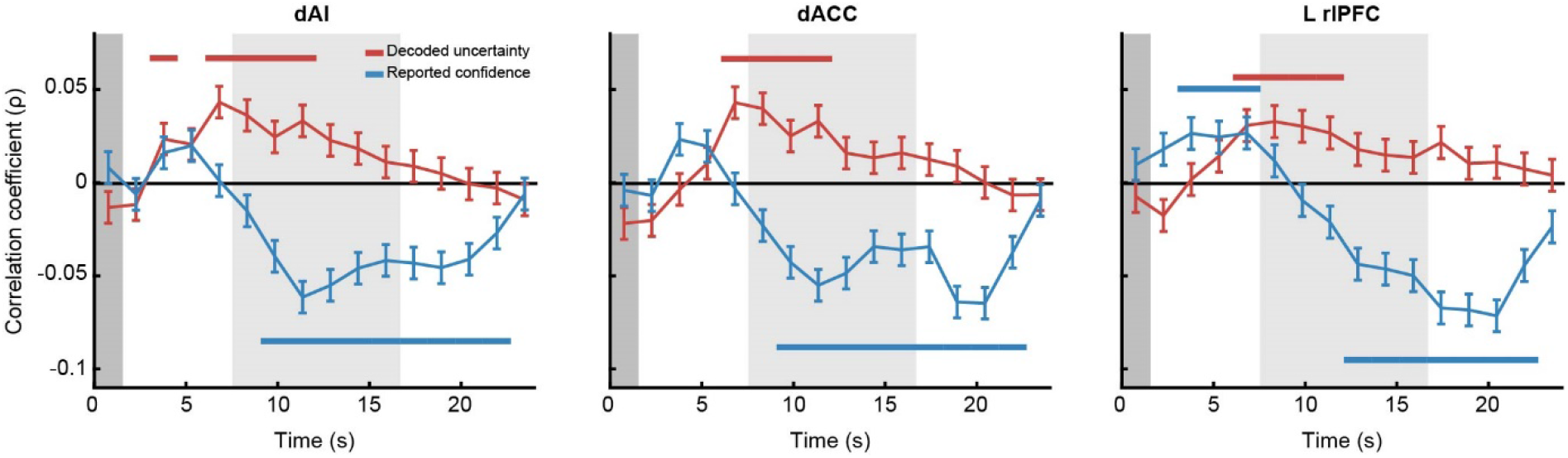
Effects of decoded uncertainty and reported confidence on the BOLD response in dAI, dACC and rlPFC, after accounting for trial-by-trial fluctuations in behavioral response times. Behavioral response time effects were linearly regressed out from decoded uncertainty and reported confidence, prior to computing the Spearman correlation coefficient between decoded uncertainty (reported confidence) and the BOLD response at different moments in time after stimulus presentation. The remaining analysis steps are identical to those in the main text. Removing the effect of behavioral response time did not qualitatively change the pattern of results in any of these ROIs. Horizontal lines indicate statistical significance (p<0.05, FWER-controlled). Dark gray area marks stimulus presentation window, light gray area denotes response window. Error bars represent ± 1 s.e.m.

**Extended Data Fig. 7.**
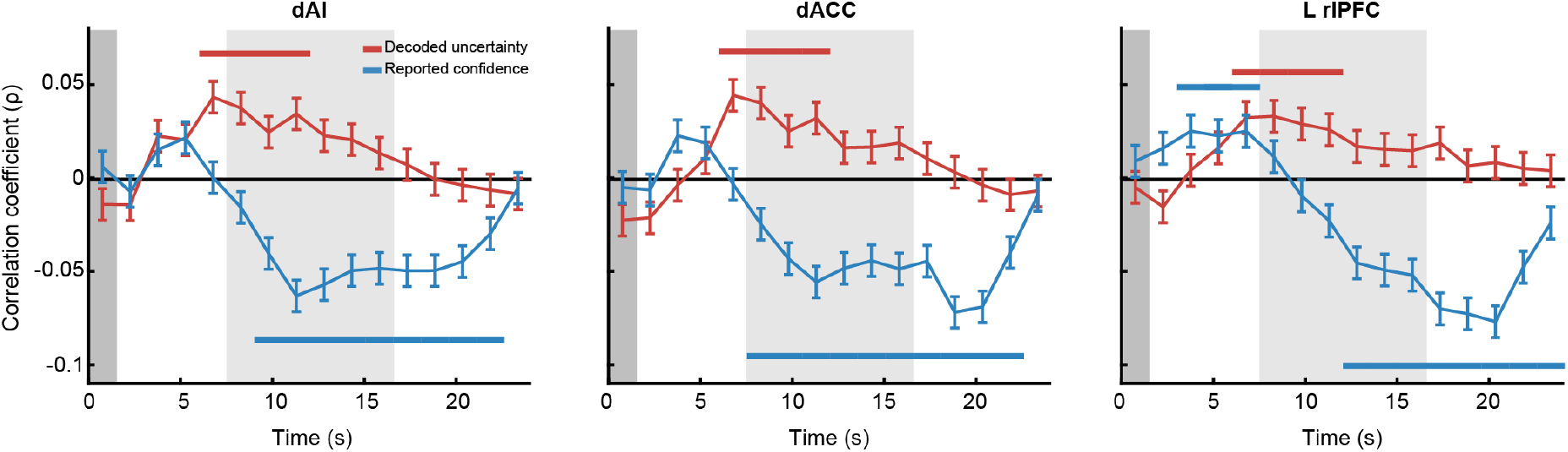
Effects of decoded uncertainty (or reported confidence) on the BOLD response in dAI, dACC and rlPFC, after controlling for confidence (or decoded uncertainty). Reported confidence (or decoded uncertainty) was linearly regressed on both decoded uncertainty (or reported confidence) and the BOLD response at different moments in time after stimulus presentation. The residuals of these fits were then used to compute the group-averaged correlation coefficient between cortical response amplitude and decoded uncertainty (red) or reported confidence (blue). For all ROIs, the results are qualitatively similar to the main results reported in Fig. 4 in the main text. Horizontal lines indicate statistical significance (p<0.05, FWER-controlled). Dark gray area marks stimulus presentation window, light gray area denotes response window. Error bars represent ± 1 s.e.m.

**Extended Data Fig. 8.**
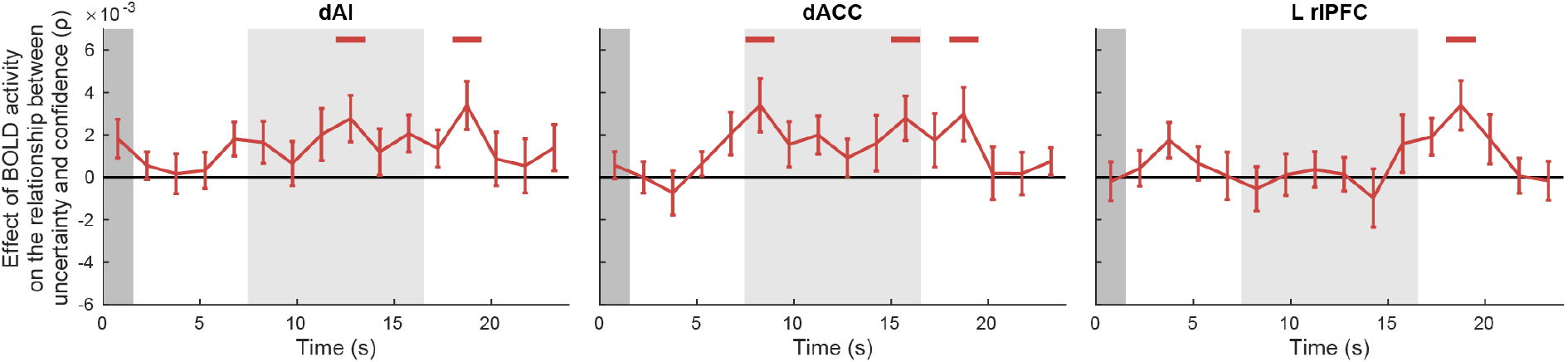
Activity in dAI, dACC, and left rlPFC mediates the relationship between decoded uncertainty and reported confidence. To assess the degree to which the cortical activity in these regions mediates the observed relationship between decoded uncertainty and reported confidence, we performed the following analysis. We first modeled both uncertainty and confidence as a function of the overall BOLD signal in a given ROI at each timepoint, and then used the residuals of these fits to compute the Spearman correlation coefficient between decoded uncertainty and reported confidence when controlled for the BOLD signal. From the resulting correlation coefficient, we subtracted the (baseline) correlation coefficient that was obtained while we did not control for the BOLD signal (see Fig. 3c). We observed a significant net effect at various moments in time, which indicates that there was a reliable reduction in the strength of the inverse (negative) correlation coefficient between uncertainty and confidence when we controlled for BOLD intensity. This suggests that the level of cortical activity in these windows (partially) mediates the relationship between decoded uncertainty and reported confidence. Horizontal lines indicate statistical significance (p<0.05, FWER-controlled). Dark gray area marks stimulus presentation window, light gray area denotes response window. Dashed lines indicate the decoding window used in the main analyses (Fig. 3b-c and Extended Data Fig. 2). Error bars represent ± 1 s.e.m.

**Extended Data Fig. 9.**
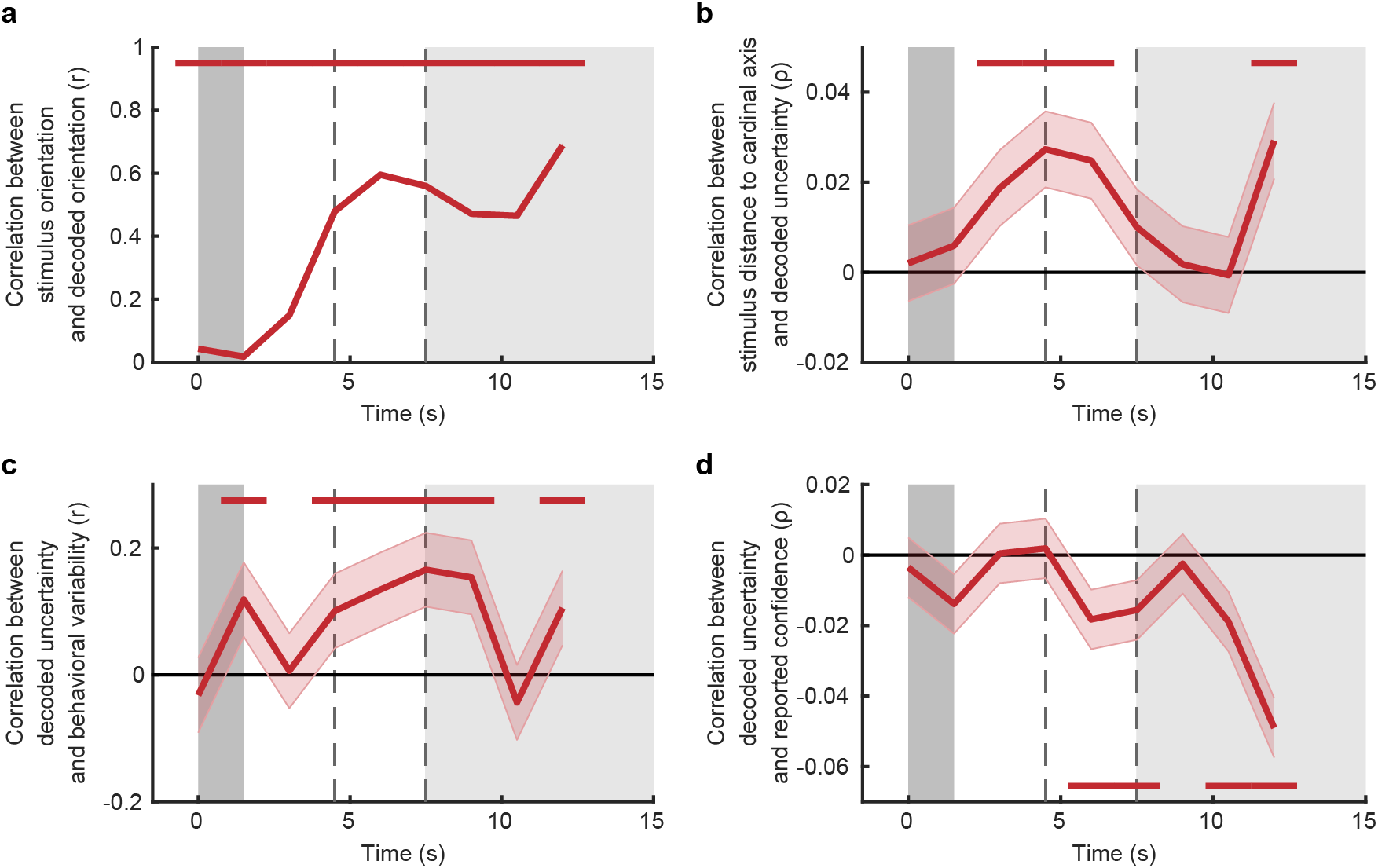
Decoding results over time. Does reported confidence similarly reflect imprecision in the cortical representation when the orientation is held in visual working memory? To address this question, the analyses of Fig. 3b-c and Extended Data Fig. 2 were repeated over time, using a sliding window of size 3 s (2 TRs). We focused on successive intervals from 1.5 s before stimulus onset to 13.5 s after stimulus onset (which roughly corresponds to the onset of the response window after accounting for hemodynamic delay). Benchmark tests verified that the decoded probability distributions reliably predict the orientation of the presented stimulus (a), and variability in the observer’s behavioral estimates (b-c) over extended periods of time. Having established that the decoded distributions meaningfully reflect the degree of imprecision in the cortical representation, we next investigated the extent to which decoded uncertainty predicts reported confidence during the retention interval. Interestingly, we found a reliable negative relationship between decoded uncertainty and reported confidence that held up well into the delay period (d). This is consistent with an imprecise working memory trace in V1-V3 that influences subjective confidence. Please note, however, that our design does not warrant strong conclusions regarding the nature of this representation: due to fMRI’s low temporal resolution, it is difficult to say whether these signals are purely perceptual or related to working memory (see e.g. ^47^, for similar rationale), and later TRs could simply reflect the visual presentation of the response bar, rather than memory-based signals. (a-d) Data are centered to the middle of the analysis window (of size 2 TRs). Horizontal lines indicate statistical significance (p<0.05, FWER-controlled). Dark gray area marks stimulus presentation window, light gray area denotes response window. Dashed lines indicate the decoding window used for the main analyses (i.e., Fig. 3b-c and Extended Data Fig. 2). Shaded regions represent ± 1 s.e.m. (standard errors in (a) are too small to be visible).

**Extended Data Fig. 10.**
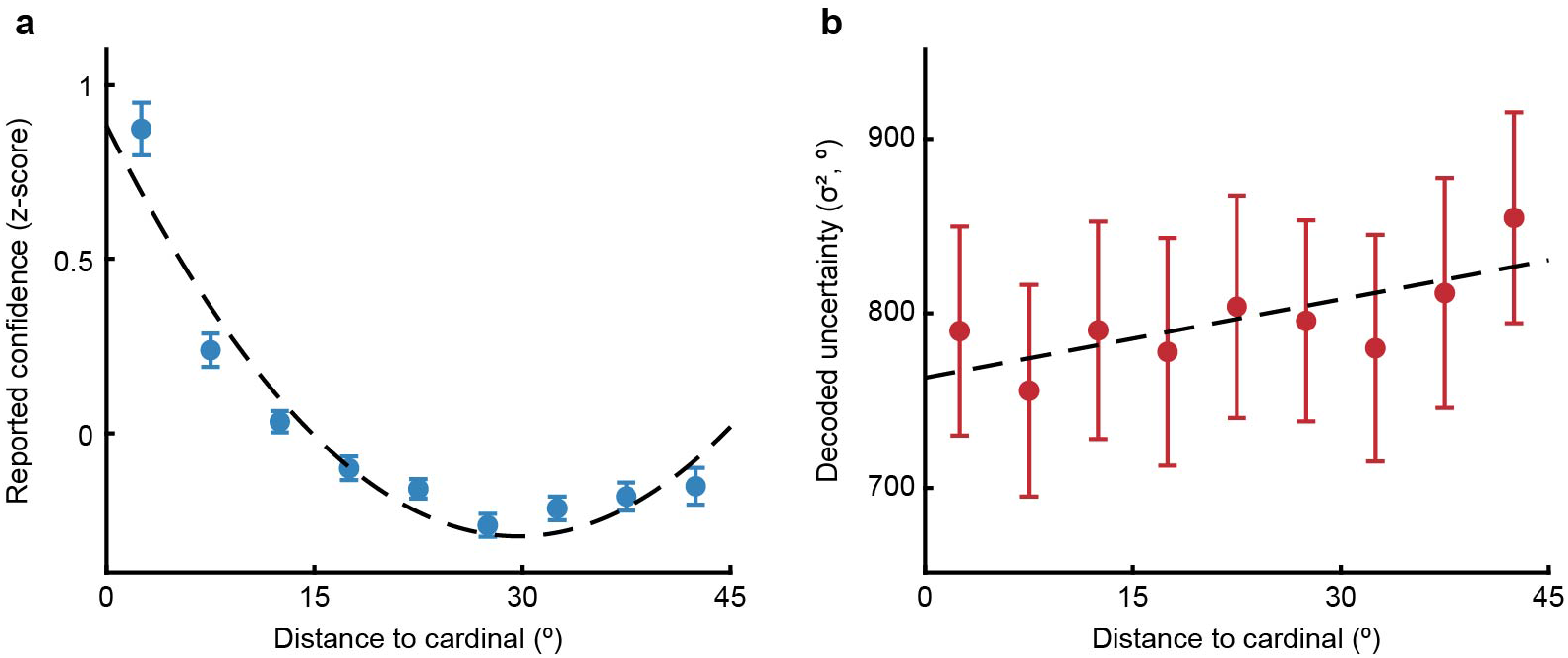
Oblique effect in reported confidence and decoded uncertainty. Effect of stimulus orientation on reported confidence (a) and decoded uncertainty (b). Each participant’s data were first binned based on the absolute distance between presented stimulus orientation and the nearest cardinal axis (equal-width bins), and then averaged across trials and finally across subjects (error bars represent ± 1 s.e.m). Dashed lines indicate best-fitting function (least-squares; quadratic for confidence, linear for decoded uncertainty). Functions were fitted on the trial-by-trial data for each participant, and averaged across participants.

## Supplementary Information

**Supplementary Table 1.** Overview of regions >5 voxels in which activity reliably co-fluctuated with decoded sensory uncertainty (whole-brain univariate analysis; p<0.05 FWER-corrected)

| Size<br>(voxels) | MNI coordinates<br>of peak (mm) |  |  | Laterality | Label |
| --- | --- | --- | --- | --- | --- |
|  | X | Y | Z |  |  |
| 5651 | -14 | -72 | -12 | bilateral | Occipital cortex / precuneus |
| 665 | -8 | 12 | 40 | bilateral | Dorsal anterior cingulate cortex (rostral cingulate zone) |
| 156 | -38 | 18 | -8 | L | Dorsal anterior insula |
| 133 | 0 | -22 | 46 | bilateral | Dorsal posterior cingulate cortex (caudal cingulate zone) |
| 82 | -4 | -16 | 28 | bilateral | Ventral posterior cingulate cortex (area 23ab) |
| 67 | -24 | 46 | 32 | L | Dorsolateral prefrontal cortex (area 9/46d, area 46) |
| 52 | -38 | 10 | 4 | L | Dorsal anterior insula |
| 41 | -12 | 12 | 66 | L | Supplementary motor area |
| 41 | 44 | 18 | 2 | R | Dorsal anterior insula |
| 26 | -6 | 38 | 18 | L | Perigenual anterior cingulate cortex (area 32d) |
| 15 | -34 | 20 | 10 | L | Dorsal anterior insula |
| 13 | -32 | -92 | -14 | L | Occipital cortex |
| 10 | 6 | 12 | 52 | R | Presupplementary motor area |
| 8 | 10 | -22 | 38 | R | Dorsal posterior cingulate cortex (caudal cingulate zone) |
| 6 | 0 | 50 | 10 | bilateral | Perigenual anterior cingulate cortex (area 32pl) |

